# Nutrient metabolism regulates insulin granule formation in the pancreatic islet β-cell via ER redox homeostasis

**DOI:** 10.1101/2021.09.22.461417

**Authors:** Kristen E. Rohli, Cierra K. Boyer, Shelby C. Bearrows, Marshall R. Moyer, Weston S. Elison, Casey J. Bauchle, Sandra E. Blom, Chi-Li Yu, Marshall R. Pope, Jianchao Zhang, Yanzhuang Wang, Samuel B. Stephens

**Author notes:** Corresponding author: Samuel B. Stephens, Ph.D., Fraternal Order of Eagles Diabetes Research Center, Department of Internal Medicine, Division of Endocrinology and Metabolism University of Iowa, Iowa City, IA 52246.

## Abstract

Defects in the pancreatic β-cell’s secretion system are well-described in Type 2 diabetes (T2D) and include impaired proinsulin processing and a deficit in mature insulin-containing secretory granules; however, the cellular mechanisms underlying these defects and the contribution of hyperglycemia to this process remain poorly understood. Here, we used an *in situ* fluorescent pulse-chase strategy and proximity labeling-based quantitative proteomics analysis to study proinsulin trafficking and demonstrate a direct link to glucose metabolism via the production of redox intermediates that facilitate proinsulin export from the ER. We show that ER export of proinsulin is delayed in T2D models resulting in decreased insulin granule formation and further demonstrate this process can be regulated by NADPH and reducing equivalent availability. Together, these data highlight a critical role for nutrient metabolism and mitochondrial dysfunction in the maladaptive remodeling of the β-cell’s secretory pathway in the decline of β-cell function in T2D.

## Introduction

In the pancreatic islet β-cell, metabolic fuel sensing functions at multiple levels to coordinate insulin secretory capacity with physiological demands of nutrient status (Boland et al., 2017). For example, ATP and other metabolic signals generated by glucose oxidation directly trigger and amplify insulin granule exocytosis through membrane depolarization and insulin granule recruitment (Gembal et al., 1992; Henquin, 2009; Henquin et al., 2002). In addition, mitochondrial metabolism of glucose and other fuels stimulates the translation of preproinsulin mRNA through a stem-loop structure in the 5’UTR, as well as ∼50 other mRNAs associated with insulin granules (Guest et al., 1989; Uchizono et al., 2007; Wicksteed et al., 2003), to ensure that β-cells meet or exceed the physiological demands of insulin release with insulin biosynthesis. Nutrient challenge can also elicit more global changes in β-cell secretory function, such as during prolonged fasting, which results in a pronounced reduction in the β-cell’s insulin secretory capacity, including insulin degranulation and increased numbers of autophagosomes and lysosomes, as part of a physiological mechanism to limit insulin release and thereby ward off hypoglycemia (Boland et al., 2018). Collectively, these observations illustrate the β-cell’s ability to use metabolic fuels as hierarchal signals to modulate secretory functions.

Loss of β-cell secretory function and insulin insufficiency have been identified as critical events in the progression of insulin resistance to Type 2 diabetes (T2D), yet the underlying cellular mechanisms are not well understood (Halban et al., 2014; Kahn et al., 2009). Overt changes in the β-cell’s secretory pathway accompany the onset of T2D and include decreased numbers of insulin-containing secretory granules, proinsulin processing deficiencies, and hyperproinsulinemia (Alarcon et al., 2016; Kahn and Halban, 1997; Like and Chick, 1970; Masini et al., 2012; Ward et al., 1984). While increased insulin exocytosis and proinsulin/insulin degradation likely contribute to the overall deficit in available insulin supply in T2D (Pasquier et al., 2019; Rhodes and Alarcon, 1994; Shrestha et al., 2020, p. 1), perturbations in the ultrastructure of the β-cell secretory organelles (ER, Golgi, autophagosomes) suggest that alterations in early stages of proinsulin sorting and trafficking also contribute to the insulin insufficiency in T2D (Alarcon et al., 2016; Like and Chick, 1970; Masini et al., 2012). Studies on early stages of proinsulin trafficking and insulin granule formation have been challenging, in part, due to the prolonged half-life of proteins within the mature secretory granule, such as insulin (t ½ ∼ 2.7 days) (Muller et al., 2017), which are also highly abundant. Nevertheless, defects in proinsulin trafficking and insulin granule formation can directly decrease insulin secretory capacity (Bearrows et al., 2019; Cao et al., 2013, p. 1; Stephens et al., 2017) and may occur in T2D (Gandasi et al., 2018); however direct measures of insulin granule formation in T2D have been lacking.

Insulin is initially synthesized as the prohormone precursor, proinsulin, which is folded via the assistance of ER resident chaperones, such as BiP (Arunagiri et al., 2018) and GRP94 (Ghiasi et al., 2019; Kim et al., 2018), while protein disulfide isomerases (PDIs), including Pdia1 and Prdx4, facilitate formation of the three critical disulfide bonds necessary for mature insulin structure (Jang et al., 2019; Tran et al., 2020; Winter et al., 2002). To maintain PDIs for sequential rounds of disulfide bond isomerization, reducing equivalents in the form of glutathione and/or thioredoxin act as the final electron donors (Birk et al., 2013; Ellgaard et al., 2018), which are recycled by NADPH-dependent enzymes, glutathione reductase and thioredoxin reductase, respectively. While the mechanisms regulating transfer of reducing equivalents from the cytosol to the ER lumen is not well understood (Cao et al., 2020; Ellgaard et al., 2018), recent studies in hepatocytes demonstrate that enhanced TCA cycle flux, which increases NADPH production and glutathione reducing equivalents (GSH), results in a more reduced ER environment (Gansemer et al., 2020). Conversely, diminished TCA metabolism and NADPH production leads to a decrease in available GSH and a more oxidized ER lumen. In β-cells, NADPH and glutathione redox fluctuate with physiological changes in glucose metabolism (Ivarsson et al., 2005; Reinbothe et al., 2009). Thus, defects in mitochondrial metabolism in T2D could directly alter β-cell ER redox poise through changes in cellular redox flux and thereby impact proinsulin folding and downstream insulin granule production. Indeed, recent work in human and rodent T2D β-cells has identified the formation of oligomeric aggregates of proinsulin arising from mis-matched intermolecular disulfide bonds (Arunagiri et al., 2019; Tran et al., 2020), suggesting that defects in proinsulin disulfide bond isomerization may be prominent in T2D. While previous studies have demonstrated a role for NADPH and redox intermediates in the regulation of insulin exocytosis (Ferdaoussi et al., 2015; Ivarsson et al., 2005; Reinbothe et al., 2009; Ronnebaum et al., 2006), the contribution of redox biology to ER function and proinsulin trafficking in the β-cell has not been investigated.

In the present study, we evaluated proinsulin trafficking and insulin granule formation in rodent and cell culture models of hyperglycemia and T2D. Using a fluorescent pulse-chase proinsulin reporter in diabetes models, we found a significant decrease in the production of insulin granules, which exhibited delayed trafficking to the plasma membrane. Using APEX2 proximity labeling to identify protein interactions with proinsulin, we found a profound increase in the enrichment of proinsulin interactions with ER oxidoreductases in β-cells cultured under hyperglycemic-like conditions, which was accompanied by diminished mitochondrial function and increased glutathione oxidation. Furthermore, we directly demonstrate a significant delay in proinsulin export from the ER in both animal and cell culture models of T2D, which can be regulated by the availability of NADPH and cellular reducing equivalents. Together, our data suggest that mitochondrial metabolism fails to maintain NADPH-derived reducing equivalents necessary to support ER function in T2D β-cells, which ultimately leads to a decline in insulin granule formation.

## Results

### Impaired proinsulin trafficking occurs in models of β-cell dysfunction

In T2D, β-cell dysfunction is accompanied by a decline in insulin-containing secretory granules (Alarcon et al., 2016; Like and Chick, 1970; Masini et al., 2012). Whether alterations to insulin granule formation contribute to this deficit is not known. To address this, we evaluated proinsulin trafficking and insulin granule formation in a rodent diabetes model. For these studies, C57BL6/J mice were placed on a Western diet (WD; 40% fat/kcal, 43% carbohydrate/kcal) or standard chow (SC) for up to 8 weeks. Increased body weight (Figure 1A) and *ad lib* fed hyperglycemia (Figure 1B) were apparent within 4 weeks of dietary intervention and persistent for the duration of diet. Reduced glucose tolerance (Figure 1C), fasting hyperinsulinemia, and impaired glucose-stimulated insulin release (Figure 1D) were observed by 8 weeks of WD. Collectively, these effects are consistent with defects in β-cell function in T2D (Halban et al., 2014; Kahn et al., 2009).

**Figure 1.**
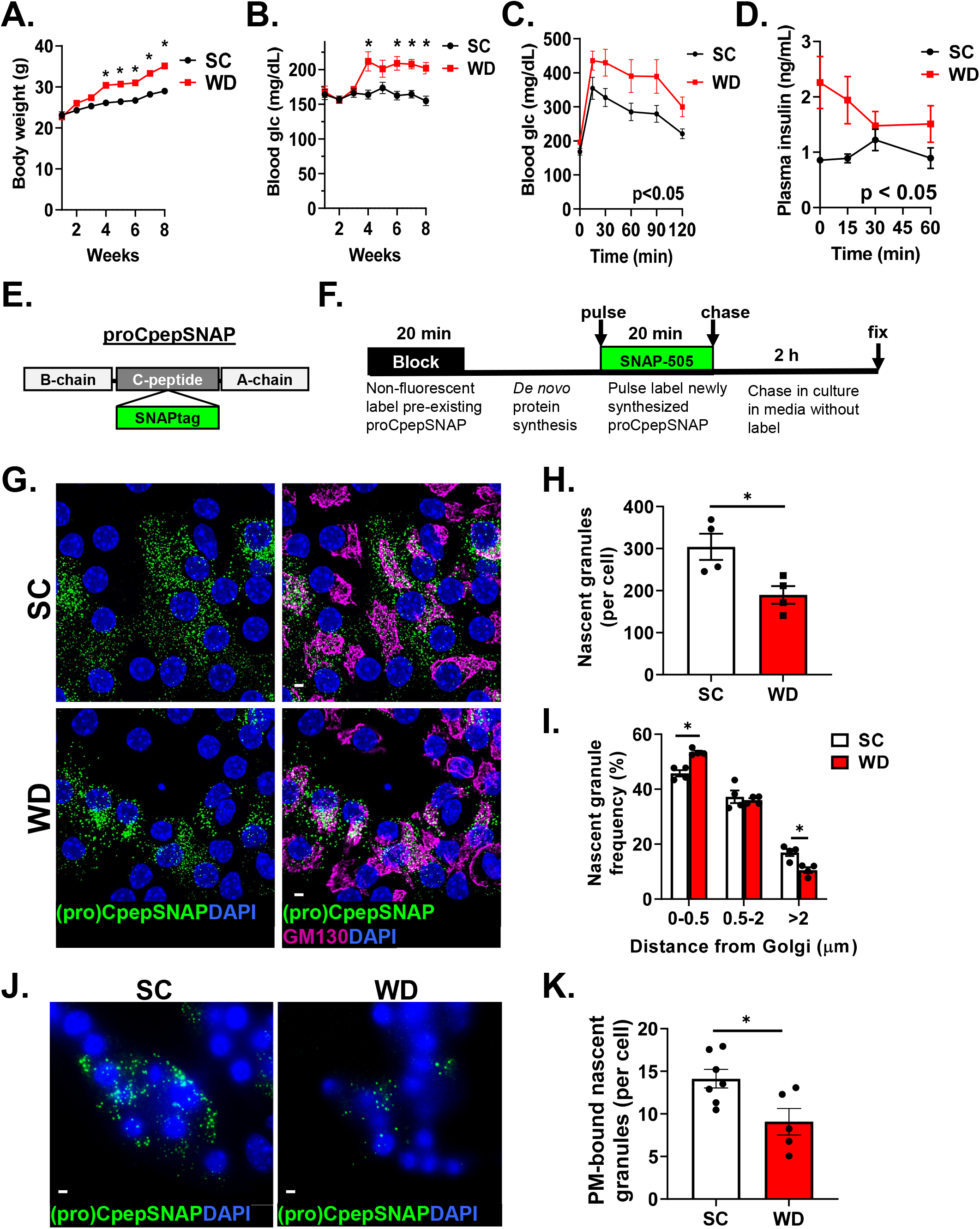
Insulin granule formation is impaired in a rodent model of islet dysfunction. 8-10 week old male C57BL6/J mice (n=5-9) were placed on a standard chow (SC) or Western diet (WD). Body weight (**A**) and *ad lib* fed blood glucose (**B**) were recorded weekly. After 8 wk of diet, 4 h fasted mice were injected i.p. with 1 mg/g bw glucose and blood glucose monitored for 2 h as indicated (**C**) and plasma sampled for insulin (**D**). (**E**) Model of the proCpepSNAP reporter with SNAPtag inserted within the C-peptide region of human preproinsulin. (**F**) Schematic of proCpepSNAP pulse-chase labeling timeline. (**G-I**) Mouse islets (8 wk, SC vs. WD C57BL6/J) were treated with AdRIP-proCpepSNAP. 48 h post-infection, islets were pulse-labeled with SNAP-505 for 20 min (green), chased for 2 h, immunostained for GM130 (magenta), and counterstained with DAPI (blue). Confocal images were collected (**G**), and the total number of proCpepSNAP-labeled nascent granules quantified (**H**) (n=4; 22-25 cells per mouse). (**I**) Frequency distribution of binned proCpepSNAP-labeled granule distances (microns) from the Golgi were quantified (n=4). proCpepSNAP pulse-chase labeling of β-cells from C57BL6/J mice (14 wk, SC or WD, n=5) were imaged by TIRF microscopy (**J**) and the total number of plasma membrane (PM)-bound proCpepSNAP-labeled granules are quantified (**K**) (n=5-7; 26-57 cells per mouse). (**G**, **J**) Scale bar = 5 μm. (**A-D**, **H**, **I**, **K**) Data represent the mean ± S.E.M. * p < 0.05 by two-way ANOVA with Sidak post-test analysis (**A-D**, **I**) or Student t test (**H**, **K**).

We previously developed an *in situ* pulse-chase fluorescent-labeling strategy based on the modified DNA repair enzyme, SNAPtag, to track nascent proinsulin transit through the secretory system and measure insulin granule formation (Bearrows et al., 2019). We inserted SNAPtag within the C-peptide region of human preproinsulin (Figure 1E), termed proCpepSNAP, and demonstrated that the trafficking, processing, and secretion of proCpepSNAP/CpepSNAP are consistent with the regulation of endogenous proinsulin/insulin (Bearrows et al., 2019). In this system, SNAP-tag labeling is a terminal event; once labeled, the reactive cysteine is not available for subsequent labeling. Taking advantage of this, proCpepSNAP expressing cells are initially pre-labeled with a non-fluorescent SNAP-tag probe to mask the existing pool of (pro)CpepSNAP (Figures 1F and S1). This is followed by a recovery time to allow *de novo* protein synthesis of proCpepSNAP, followed by a pulse of fluorescent (SNAP-505) labeling. A subsequent 2 h chase in media allows our analysis to focus on Golgi exit and insulin granule formation (Bearrows et al., 2019). In the present study, islets were isolated from SC and WD fed mice and treated with recombinant adenovirus expressing proCpepSNAP under control of the rat insulin promoter (RIP) to ensure β-cell specific expression. Following pulse-chase labeling, we observed the appearance of numerous nascent proinsulin (proCpepSNAP)-containing granules throughout the cell body in primary β-cells from non-diabetic, SC fed animals (Figure 1G). In contrast, β-cells from WD fed animals displayed a substantial reduction in the total number of nascent, proCpepSNAP-labeled granules formed (Figure 1G, H). Moreover, distance quantitation of these data revealed a significant increase in the frequency of nascent proinsulin granules within 0.5 µm of the Golgi (GM130 marker) and a decrease in granules greater than 2 µm from the Golgi in β-cells from WD compared to SC fed mice (Figure 1I). Furthermore, analysis by TIRF microscopy demonstrated a substantial decrease in the number of nascent (proCpepSNAP-labeled) granules trafficking to the plasma membrane (≤ 150 nm) in the WD model (Figure 1J, K). Together, these data formally demonstrate an impairment in insulin granule formation and trafficking in a rodent model of diet-induced diabetes, consistent with past reports of reduced numbers of mature insulin granules in T2D (Like and Chick, 1970; Masini et al., 2012).

To further investigate alterations to insulin granule formation, we examined islets isolated from overtly diabetic C57BLKS/J-db/db mice compared to normoglycemic C57BLKS/J-db/+ controls, which displayed a profound impairment in glucose tolerance (Figure 2A) accompanied by the loss of glucose-regulated insulin secretion (Figure 2B). In this model, we observed significant decreases in the expression of granule markers, PC2 and CPE, in islets from diabetic db/db mice compared to non-diabetic db/+ controls (Figure 2C, D), consistent with previous reports (Kang et al., 2019). Next, we examined an insulinoma cell culture model of β-cell dysfunction. In these studies, 832/3 cells were cultured with either BSA alone or an oleate/palmitate mixture conjugated to BSA (1 mM; OP) at either normal (7.5 mM) or elevated (20 mM) glucose. Following 72 h culture, we measured insulin secretion after 1 h static incubation at basal (2.5 mM) and stimulatory (12 mM) glucose. We observed no effect of fatty acid treatment alone (OP) on β-cell function, whereas elevated glucose culture alone resulted in increased basal insulin secretion. In contrast, oleate/palmitate and elevated glucose together (OPG) resulted in both increased basal insulin secretion and impaired glucose-stimulated insulin secretion (Figure 2E). By immunoblot, we observed a decrease in the expression of granule markers, PC2 and CPE, in OPG-cultured 832/3 cells compared to BSA controls, whereas another key granule protein, CgB, was unaffected (Figure 2F, G). Using our fluorescent pulse-chase proCpepSNAP reporter system, we demonstrate that OPG-cultured 832/3 cells displayed increased retention of nascent proinsulin-containing (proCpepSNAP-labeled) granules proximal to the Golgi (less than 3 µm) and decreased numbers of granules distal to the Golgi (greater than 5 µm; Figure 2H, I), similar to β-cells from WD mice (Figure 1I). Additional analysis by TIRF microscopy confirmed the decrease in nascent insulin granules reaching the plasma membrane after a 4 h chase in 832/3 cells cultured under high glucose (HG; 20 mM) compared to normal glucose (NG; 7.5 mM) conditions (Figure 2J, K). Following a prolonged chase (> 18 h), we observed a similar number of nascent granules (proCpepSNAP-labeled) in the BSA (control) and OPG-cultured cells (Figure 2L, M), suggesting that defects in insulin granule formation in our OPG model stemmed from trafficking delays, rather than reduced expression and/or stability of our reporter. Last, we examined total insulin granules by density sedimentation in control (BSA) and OPG-cultured 832/3 cells. Here, we observed a significant shift of insulin-containing granules to lighter density fractions from OPG cultured cells, suggesting there was an alteration to granule composition (Figure 2N). Collectively, these data demonstrate impairments in insulin granule formation and alterations to granule composition in both cell culture and animal models of β-cell dysfunction.

**Figure 2.**
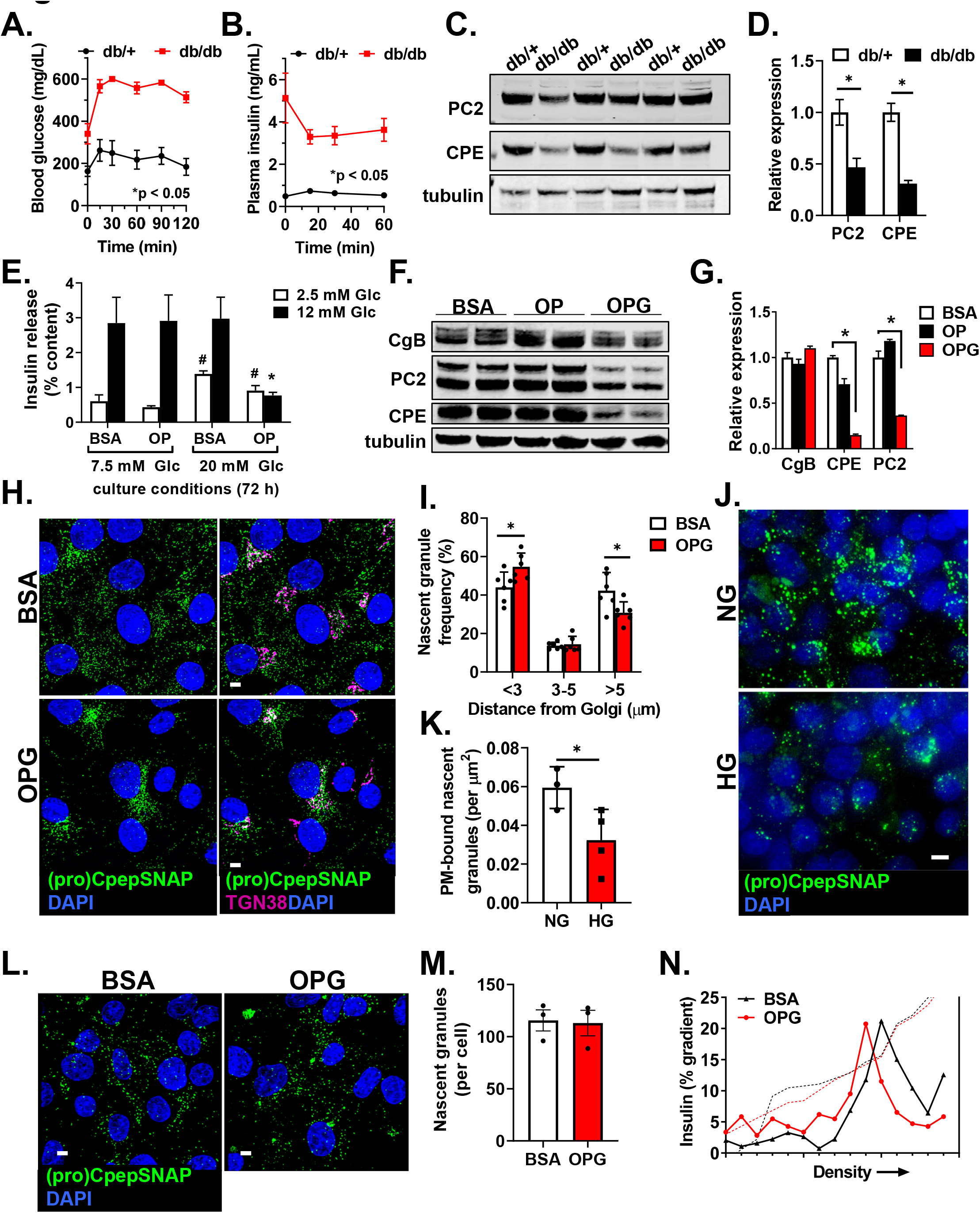
Elevated glucose and fatty acid culture impair β-cell function and proinsulin trafficking. (**A**-**B**) 10-14 week old C57BLKS/J db/+ and db/db mice (n=4-5 per group) were fasted for 4 h and injected i.p. with 1mg/g bw glucose. Blood glucose was monitored for 2 h as indicated (**A**) and plasma was sampled for insulin (**B**). Islet lysates from 10-14 wk old C57BLKS/J db/+ vs. db/db mice were analyzed by immunoblots (**C**) and relative protein expression quantified following normalization to the tubulin loading control (**D**). (**E**-**N**) 832/3 cells were cultured for 72 h in control media supplemented with BSA, media containing oleate:palmitate (2:1, 1mM OP), or media containing oleate/palmitate (2:1, 1 mM) and elevated glucose (20mM; OPG) as indicated. (**E**) Glucose-stimulated insulin secretion was measured by static incubation in media containing 2.5 mM Glc followed by 12 mM Glc for 1 h each. Immunoblots from whole-cell lysates were probed with CgB, PC2, and CPE antibodies are shown (**F**) and relative protein expression was quantified following normalization to the tubulin loading control (**G**) (n=3). 832/3 cells stably expressing proCpepSNAP were pulse-labeled with SNAP-505 for 20 min (green), chased for 2 h (**H**, **I**), 4 h (**J**, **K**), or 18-20 h (**L**, **M**), immunostained for TGN38 (magenta) and counterstained with DAPI (blue). Representative images of 2 h chase are shown (**H**) and frequency distribution of binned proCpepSNAP-labeled nascent granule distance from the Golgi determined (**I**) (n=6; 35-37 cells per group; scale bar = 5 μm). The total number of nascent (proCpepSNAP-labeled) insulin granules bound to the plasma membrane (PM) via TIRF microscopy after a 4 h chase is shown (**J**) and normalized to cell area (**K**) (n=3; 4 images per condition). The total number of nascent (proCpepSNAP-labeled) insulin granules following a 18 h chase is shown (**L**) and quantified (**M**) (61-63 cells per condition, n=3). (**N**) Cell lysates were resolved on 8-23% iodixanol density gradients. Solid line represents insulin as determined by ELISA from gradient fractions; dotted line represents gradient density (absorbance at 340 nm). (**A**, **B**, **D**, **E**, **G**, **I**, **K**, **M**) Data represent the mean ± S.E.M. * p < 0.05 by repeated measures ANOVA with Sidak post-test analysis (**A**, **B**), Student t test (**D**, **K**, **M**), or two-way ANOVA with Sidak post-test analysis (**E**, **G**, **I**). (**E**) * comparison of secretion response at stimulatory glucose to control (BSA, 7.5 mM); # comparison of secretion response at basal glucose to control (BSA, 7.5 mM).

### Proximity labeling of the proinsulin interactome reveals an ER export delay

We hypothesized that changes in the proteins interacting with proinsulin along the secretory pathway contribute to the decrease in formation of insulin secretory granules. To address this, we used an *in situ* proximity labeling strategy to systematically map the proinsulin interactome via the engineered ascorbate peroxidase, APEX2, inserted within the C-peptide region of human preproinsulin (proCpepAPEX2), which is analogous to our proCpepSNAP construct. In the presence of hydrogen peroxide, APEX2 generates a short-lived (<1 ms), membrane impermeable, biotin-phenoxy radical that biotinylates neighboring proteins *in situ* within a small labeling radius (20 nm; labeling time < 1 min) allowing us to capture transient protein interactions that are difficult to monitor with traditional approaches (Hung et al., 2016; Lam et al., 2015). Initial validation studies demonstrated that the expression of V5-tagged proCpepAPEX2 in 832/3 cells co-localized with insulin by immunostaining (Figure 3A). Furthermore, we observed substrate-dependent (biotin-phenol and hydrogen peroxide) biotinylation of putative proinsulin-interacting proteins identified with a fluorescent streptavidin probe (Figure 3B; endogenous biotin-containing carboxylases are detected independent of APEX2 labeling). Affinity purification of proCpepAPEX2 biotinylated proteins (Figure 3C) revealed interactions with the ER resident chaperone, GRP94, and granin protein, CgB, (Figure S2A) which have both been demonstrated to regulate proinsulin trafficking (Bearrows et al., 2019; Ghiasi et al., 2019; Kim et al., 2018). In contrast, no labeling of the plasma membrane SNARE, syntaxin-4, was observed, as expected (Figure S2A).

**Figure 3.**
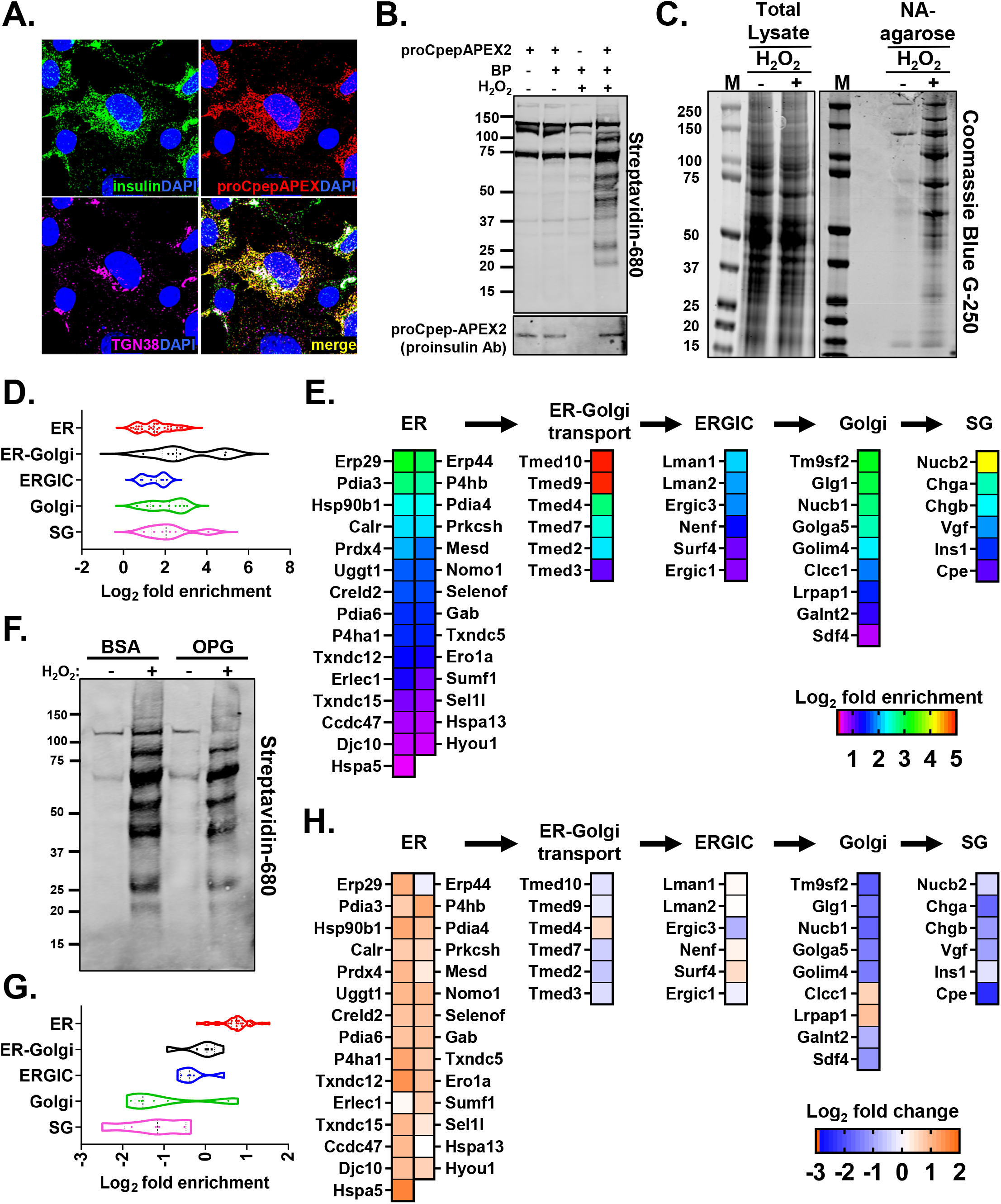
APEX2 labeling reveals increased proinsulin interactions with ER resident proteins in 832/3 cells cultured with elevated glucose and fatty acids. (**A**) 832/3 cells expressing proCpepAPEX2 were immunostained for insulin (green), V5 (proCpepV5APEX2, red), TGN38 (magenta), and counterstained with DAPI (blue). Scale bar = 5 μm. (**B**) proCpepAPEX2 expressing 832/3 cells were treated with biotin-phenol (BP) and H_2_O_2_ as indicated. Cell lysates were analyzed by Western blot probed with fluorescent streptavidin or proinsulin antibody. (**C**) Aliquots of whole cell lysates compared to Neutravidin (NA) agarose resin-enriched fractions were resolved by LDS-NuPAGE and analyzed by Coomassie blue staining. (**D, E**) Neutravidin agarose purified proteins from proCpepAPEX2 expressing 832/3 cells treated with or without H_2_O_2_ were identified by quantitative proteomics (n=4). Proteins enriched 1.2-fold or greater in peroxide-treated compared to non-peroxide-treated samples are displayed by mean-log_2_-fold enrichment as a function of subcellular location (**D**) or as a heatmap to visualize the changes of individual proteins (**E**). (**F-H**) 832/3 cells stably expressing proCpepAPEX2 were cultured for 72 h in control media supplemented with BSA or media containing oleate/palmitate (2:1, 1 mM) and elevated glucose (20 mM; OPG) as indicated. Cell lysates were analyzed by Western blot probed with fluorescent streptavidin (**F**). (**G, H**) Neutravidin agarose purified proteins were identified by quantitative proteomics. Proteins enriched 1.2-fold or greater in BSA (control) cultured cells compared to non-peroxide treated samples were further analyzed as the mean (n=4) of the relative fold (log_2_) change of OPG compared to BSA cultured cells. Data are displayed as a function of subcellular location for individual proteins (**G**) and as a heatmap (**H**).

Using the proximity labeling system, we identified proinsulin-interacting proteins by quantitative proteomics. We first compared peroxide (APEX2 dependent labeling) to non-peroxide (APEX2 independent labeling) treatments (Figure 3C) and identified 74 proteins with greater than 1.2-fold enrichment, which we categorized by their subcellular locations (Figure 3D, E). As expected, numerous ER resident chaperones (Hspa5, Hsp90b1), ER oxidoreductases (Pdia1/P4hb, Prdx4), and secretory granule proteins (CgB, Vgf) were observed to interact with proinsulin, which have all been previously demonstrated to regulate proinsulin folding (Arunagiri et al., 2018; Ghiasi et al., 2019; Jang et al., 2019; Kim et al., 2018; Tran et al., 2020; Winter et al., 2002) and trafficking (Bearrows et al., 2019; Stephens et al., 2017). We next examined changes in the proinsulin interactome in 832/3 cells cultured with elevated glucose and fatty acids (OPG) compared to control (BSA) cells (Figure 3F). Analysis by subcellular location revealed a selective increase in proinsulin interactions with ER resident proteins and a corresponding decrease in interactions with proteins residing in more distal secretory compartments (Golgi, secretory granule) (Figure 3G, H). Enriched interactions with the majority of ER proteins, such as BiP (Hspa5) and GRP94 (Hsp90b1), occurred independent of changes in gene expression (Figure 4A-C), whereas a subset of ER oxidoreductases, including Txndc12 and Prdx4, were modestly upregulated (Figure 4C). Similarly, decreased interactions with various secretory proteins, such as TMED9, TMED10, and CgB, were not accompanied by changes in protein expression (Figure 4A, B and 2F, G).

**Figure 4.**
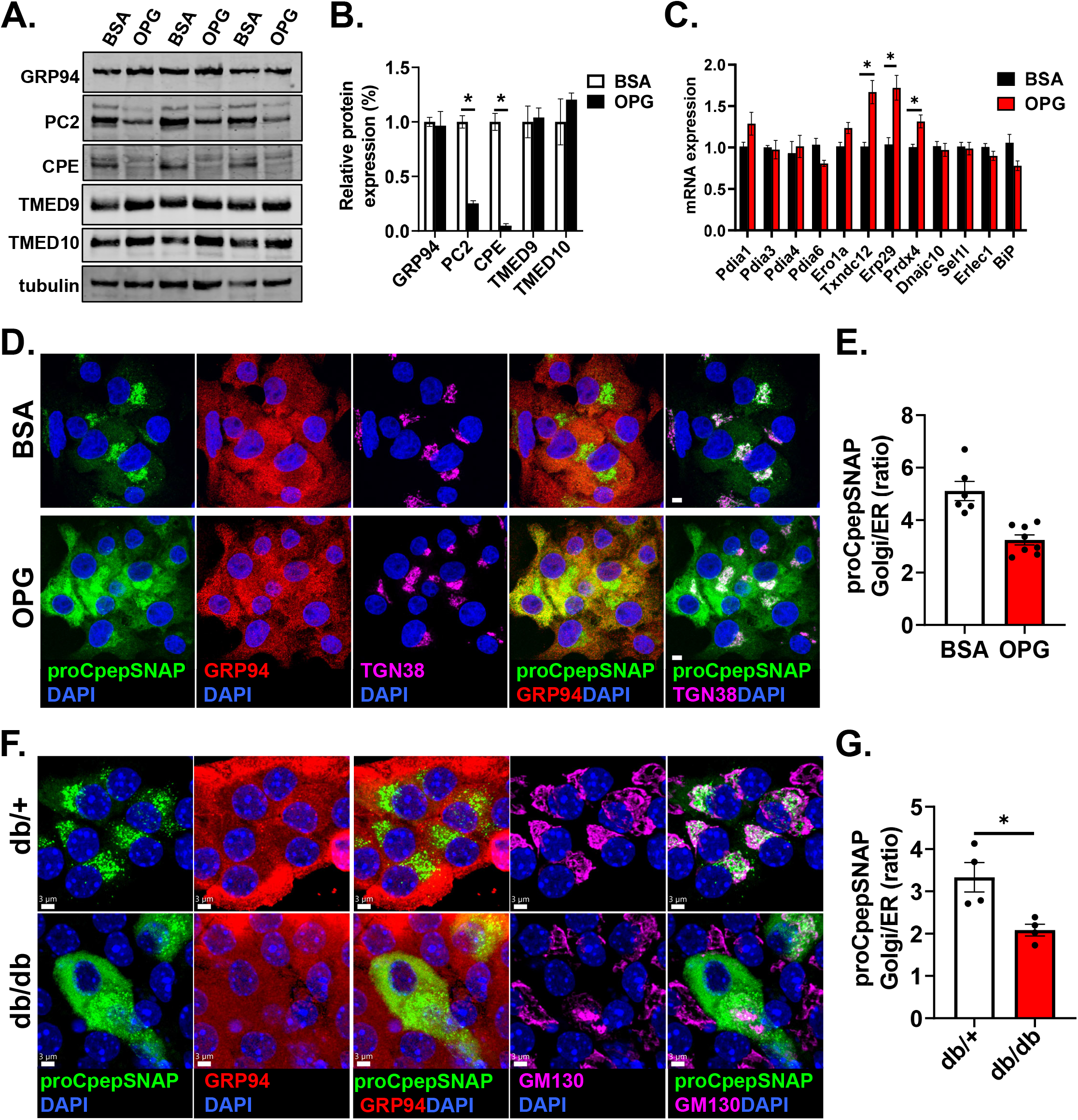
Delayed ER export of proinsulin in animal and cell culture models of hyperglycemia. (**A**-**E**) 832/3 cells were cultured for up to 72 h in control media supplemented with BSA or media containing oleate/palmitate (2:1, 1 mM) and elevated glucose (20 mM; OPG) as indicated. Cell lysates were analyzed by immunoblot (**A**) and quantified following normalization to the tubulin loading control (**B**) (n=3). (**C**) mRNA expression levels were determined by qRT-PCR (n=3). (**D-E**) 832/3 cells stably expressing proCpepSNAP were pulse-labeled with SNAP-505 for 20 min (green) then fixed and immunostained for GRP94 (red), TGN38 (magenta), and counterstained with DAPI (blue). Representative images (**D**) are shown (scale bar = 5 μm) and the ratio of proCpepSNAP fluorescence coincident with the Golgi compared to ER quantified (**E**) (n=6-8; 48-64 cells). (**F-G**) Isolated mouse islets (C57BLKS/J db/+ vs. db/db) were treated with AdRIP-proCpepSNAP. 48 h post-infection, islets were pulse-labeled with SNAP-505 for 20 min (green), chased for 10 min, immunostained for GM130 (magenta), and counterstained with DAPI (blue). Representative images (**F**) are shown (scale bar = 3 μm) and the ratio of proCpepSNAP fluorescence coincident with the Golgi compared to ER quantified (**G**) (n=4; 6-36 cells). (**B**, **C**, **E**, **G**) Data represent the mean ± S.E.M. * p < 0.05 by Student t test (**B**, **C**, **E**, **G**).

Based on our observation of increased interactions of proinsulin with ER proteins in a model of β-cell dysfunction, we hypothesized that ER export of proinsulin is delayed, which leads to impairments in insulin granule formation. To test this, we modified our pulse-chase labeling protocol to capture early-stage transit of proinsulin from the ER to Golgi (Figure S1B) using GRP94 and TGN38 immunostaining to demarcate the ER and Golgi, respectively. As expected, examination of control (BSA-cultured) cells demonstrated successful trafficking of newly synthesized proinsulin (proCpepSNAP) to the Golgi, which was largely absent from the ER (Figure 4D). In contrast, pulse-chase labeling in OPG-cultured β-cells revealed a substantial retention of nascent proinsulin coincident with the ER and very little detection within the Golgi. Quantitation revealed a significant decrease in the ratio of Golgi to ER proCpepSNAP fluorescence intensity in OPG-cultured cells (Figure 4E). We also examined ER-Golgi proinsulin trafficking in β-cells from overtly diabetic C57BLKS/J-db/db mice. While β-cells from normoglycemic db/+ controls displayed the expected trafficking of newly synthesized proinsulin (proCpepSNAP) from the ER (GRP94) to the Golgi (GM130), we observed a substantial retention of nascent proinsulin within the ER in β-cells from db/db diabetic mice (Figure 4F) with a significant decrease in the Golgi to ER proCpepSNAP fluorescence (Figure 4G).

The accumulation of proinsulin within the ER could stem from activation of the ER stress response, which has been suggested to contribute to the development of β-cell dysfunction in T2D (Yong et al., 2021). While expansion of ER membranes has been demonstrated in both human and rodent T2D β-cells (Alarcon et al., 2016; Boland et al., 2017; Marchetti et al., 2007; Masini et al., 2012); dilation of the ER lumen, which is a key hallmark of proteotoxic stress, is not commonly observed. Consistent with these latter data, the increased retention of nascent proinsulin in the ER was not accompanied by distension of the ER lumen or other changes in ER ultrastructure in β-cells from diabetic db/db mice (Figure 5A). In 832/3 cells cultured with fatty acid alone (OP) or fatty acid plus elevated glucose (OPG), we failed to detect upregulation of ER stress markers, ATF4, CHOP, XBP-1(s/u), GADD34 or BiP by qRT-PCR as compared to thapsigargin-treated cells (Figure 5B). Similarly, expression of CHOP and cleaved caspase 3 by immunoblot were not detected under OPG culture conditions (Figure S3A). Furthermore, pulse-chase analysis using our fluorescent proCpepSNAP reporter revealed that the chemical chaperone, TUDCA, which has been used to ameliorate ER stress (Ozcan et al., 2006), could not rescue the proinsulin ER export delay observed in OPG-cultured 832/3 cells (Figure S3B).

**Figure 5.**
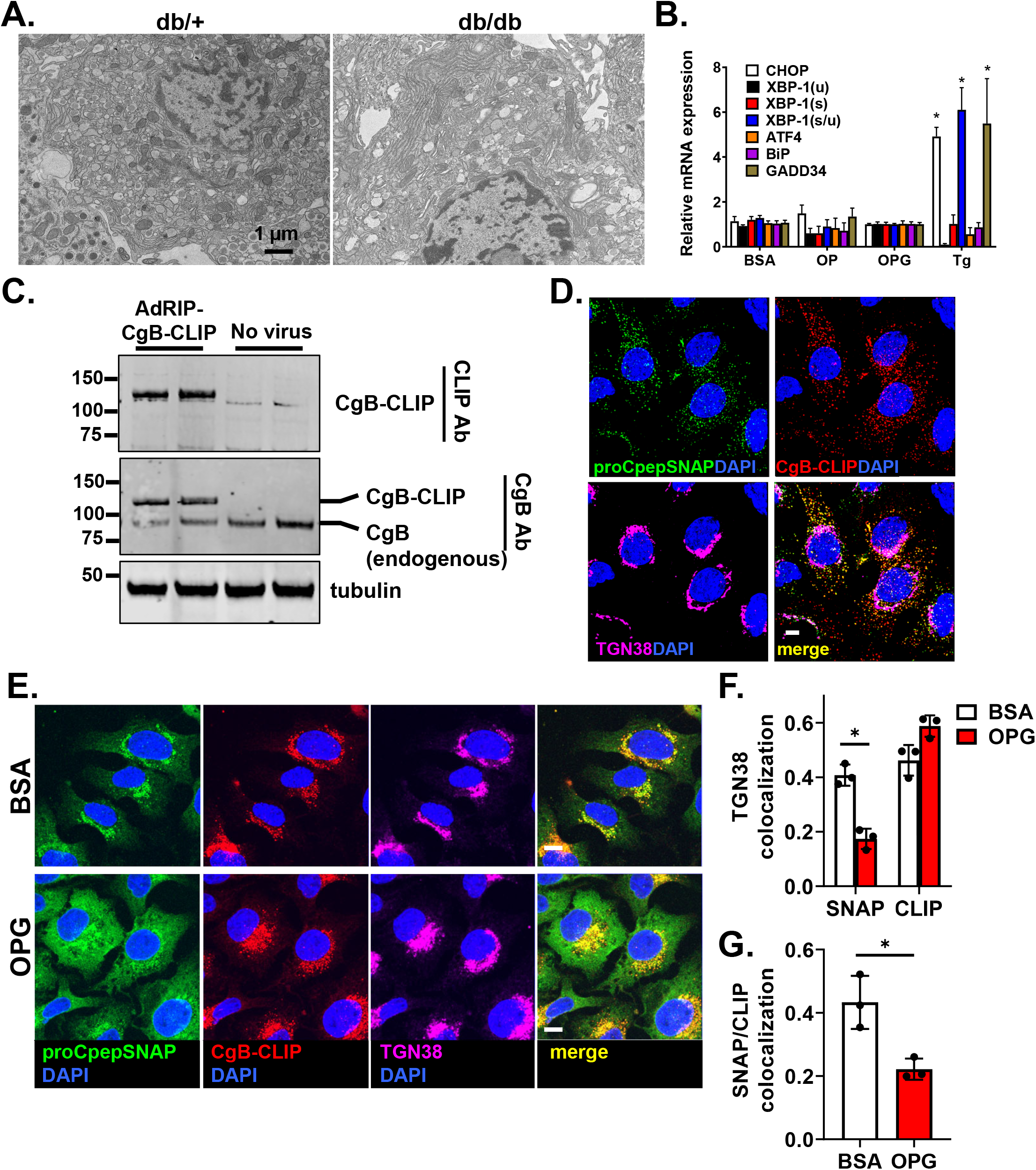
Delayed ER export delay of proinsulin is specific and independent of overt ER stress. (**A**) Islets isolated from C57BLKS/J db/+ vs. db/db (n=3) were examined for ultrastructure by electron microscopy and representative micrographs depicting the ER are shown. (**B**, **E**-**G**) 832/3 cells were cultured for 72 h in control media supplemented with BSA, media containing oleate:palmitate (2:1, 1 mM OP) or media containing oleate:palmitate (2:1, 1 mM) and elevated glucose (20 mM; OPG) as indicated. (**B**) mRNA expression was examined by qRT-PCR (n=4-6). Cells were treated for 18 h with thapsigargin (50 nM) as indicated. (**C**) 832/3 cells treated with AdRIP-CgB-CLIP were analyzed by immunoblot with antibodies raised against CLIP or endogenous CgB as indicated. (**D**) 832/3 cells expressing proCpepSNAP and CgB-CLIP were pulse-labeled with SNAP-505 (green) and CLIP-TMR (red) for 20 min, chased for 2 h, prior to fixation. Cells were immunostained for TGN38 (magenta), and counterstained with DAPI (blue). Representative images are shown (scale bar = 5 μm). (**E-G**) 832/3 cells expressing proCpepSNAP and CgB-CLIP were pulse-labeled with SNAP-505 (green) and CLIP-TMR (red) for 20 min, fixed and immunostained for TGN38 (magenta), and counterstained with DAPI (blue). (**E**) Representative images are shown (scale bar = 5 μm). Mander’s correlation coefficient (MCC) was used to determine colocalization of labeled proCpepSNAP (SNAP) vs CgB-CLIP (CLIP) with TGN38 (**F**) or proCpepSNAP (SNAP) with CgB-CLIP (CLIP) as indicated (**G**) (n=3; 53-70 cells per condition). (**B**, **F**, **G**) Data represent the mean ± S.E.M. * p < 0.05 by two-way ANOVA with Sidak post-test analysis (**B**, **F**) or Student t test (**G**).

To examine if the ER export delay was specific to proinsulin or instead was a general defect in ER protein export, we examined ER-Golgi trafficking of another granule protein, CgB, using an analogous pulse-chase system, CLIPtag. Notably, CLIPtag uses a distinct substrate from SNAPtag allowing us to perform dual pulse-chase studies of both proteins in the same cell (Gautier et al., 2008). We generated a C-terminal CLIP-tagged CgB reporter, which was expressed at levels similar to endogenous CgB (Figure 5C) and co-trafficked with proCpepSNAP into punctate structures following a 2 h pulse-chase (Fig 5D), likely representing maturing insulin granules. Using this dual pulse-chase system to examine ER-Golgi trafficking (Figure S1), we demonstrate that in BSA-cultured (control) 832/3 cells, nascent proinsulin (proCpepSNAP) and CgB (CgB-CLIP) similarly trafficked to the Golgi following pulse-chase labeling (Figure 5E, F). In OPG-cultured 832/3 cells, CgB-CLIP trafficking from the ER to Golgi occurred without delay and mirrored control (BSA) cells (Figure 5E) with similar co-localization to the Golgi marker, TGN38 (Figure 5F). This was in stark contrast to proinsulin, which was retained in the ER in OPG-cultured 832/3 cells (Figure 5E), displaying reduced co-localization with both CgB-CLIP (Figure 5G) and the Golgi marker TGN38 (Figure 5F). Collectively, these data demonstrate a specific delay in ER export of proinsulin accompanying chronic exposure to hyperglycemic and hyperlipidemic-like conditions, which is independent of overt ER stress or a general impairment in ER protein export.

### Metabolic regulation of ER redox homeostasis

In hepatocytes, mitochondrial activity can directly influence ER redox homeostasis via the generation of glutathione (GSH) reducing equivalents (Gansemer et al., 2020). Whether a similar link exists in β-cells is not known, but alterations in ER redox poise could explain the defects in ER export of proinsulin in diabetes models. To test this, we first examined mitochondrial function in control (BSA) and OPG-cultured 832/3 cells and demonstrate a significant decrease in overall mitochondrial oxidative consumption (Figure 6A), including a decrease in basal respiration (Figure S4A), maximum respiration (Figure S4B), and ATP production (Figure S4C), consistent with past reports (Haythorne et al., 2019). Furthermore, decreased mitochondrial function coincided with a corresponding shift in the redox balance of glutathione to the oxidized state (GSSG; Figure 6B) that was not simply due to an increase in the production of reactive oxygen species (Figure 6C). Because reduced glutathione (GSH) can be a final electron donor for PDIs regulating disulfide bond isomerization (Birk et al., 2013; Ellgaard et al., 2018), we reasoned that the decrease in GSH may limit proinsulin disulfide bond formation leading to an ER export delay as previously suggested (Arunagiri et al., 2019; Tran et al., 2020). To test this, we examined if addition of chemical reducing equivalents, DTT, could rescue the proinsulin trafficking delay in OPG-cultured β-cells. Note that the concentration of DTT (0.5 mM) used did not impact β-cell function following overnight exposure (Figure S4D), but did lower the GSSG/GSH ratio (Figure S4E). In control (BSA cultured) cells, acute addition of DTT (4 h) had no discernible impact on ER to Golgi trafficking of newly synthesized, proCpepSNAP-labeled, proinsulin as compared to the vehicle control (Figures 6D, E and S4F). In contrast, DTT treatment restored ER to Golgi proinsulin trafficking in OPG-cultured β-cells. These data suggest that OPG culture can lead to an insufficient supply of reducing equivalents necessary to maintain efficient ER export of proinsulin (Figure 6F).

**Figure 6.**
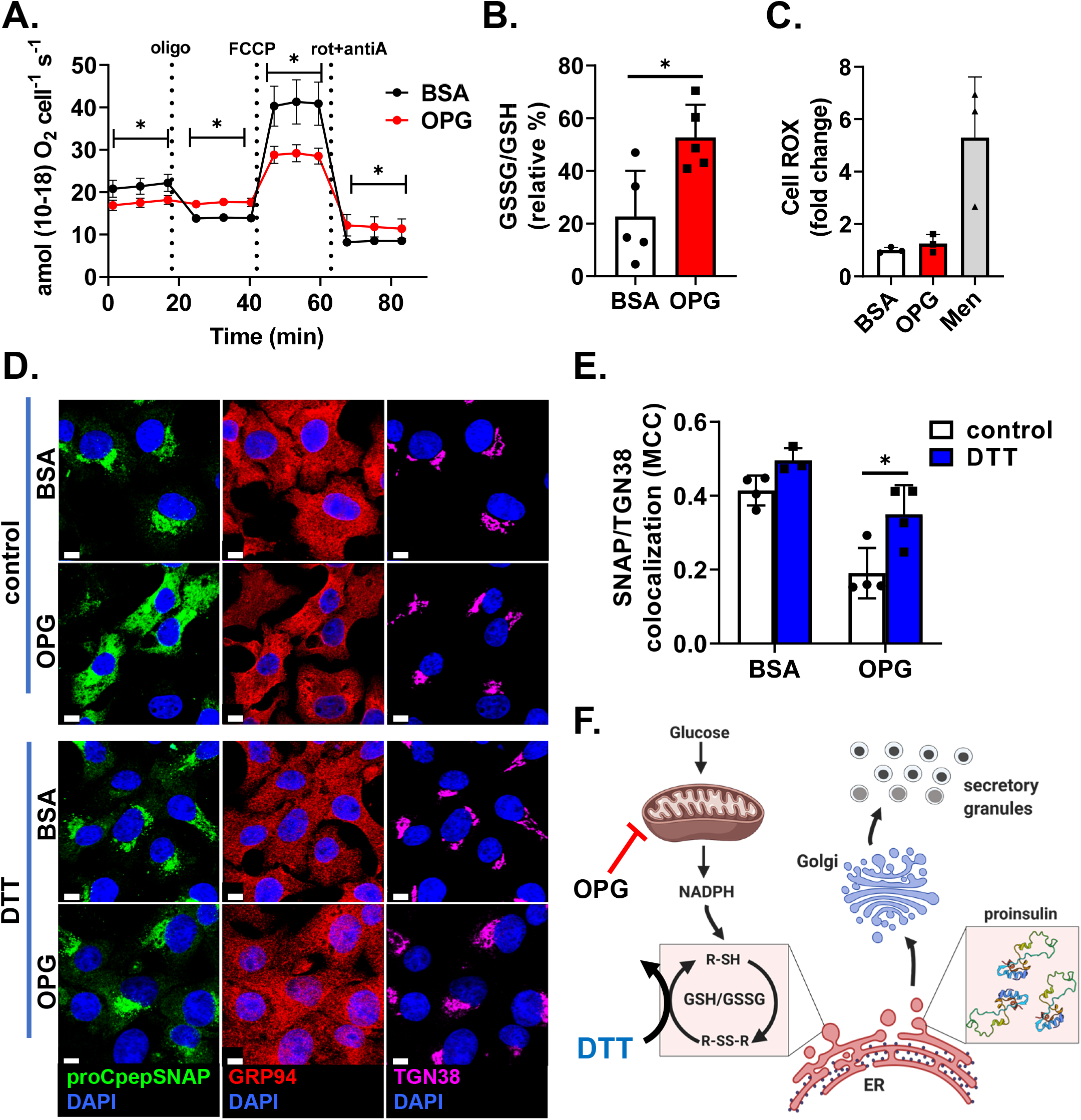
Proinsulin ER export delay can be rescued by chemical reducing agent. 832/3 cells were cultured for 72 h in control media supplemented with BSA (1%) or media containing oleate/palmitate (2:1, 1 mM) and elevated glucose (20 mM; OPG) as indicated. (**A**) Oxygen consumption rate (OCR) was measured at stimulatory glucose (20 mM) and following sequential addition of oligomycin (2.5 µM), FCCP (1.25 µM), and rotenone plus antimycin A (2 µM) as indicated (n=3 independent experiments). (**B**) 832/3 cells treated with AdRIP-Grx1-roGFP2 were imaged at 470/535 nm and 395/510 nm and re-imaged following DTT (10 mM) and diamide (5 mM) treatment for 12 minutes each (n=5). Ratiometric intensities were normalized to DTT (0%) and diamide (100%). (**C**) Cell ROX staining was used to measure ROS with 1 h menadione treatment (Men; 100 μM) as a positive control (n=3). (**D, E**) 832/3 cells stably expressing proCpepSNAP were treated with vehicle (control) or DTT (0.5 mM) for 4 h prior to being pulse-labeled with SNAP-505 for 20 min (green), fixed, and immunostained for TGN38 (magenta), and counterstained with DAPI (blue). Representative images (**D**) are shown (scale bar = 5 μm) and Mander’s correlation coefficient (**E**) was used to determine the colocalization of labeled proCpepSNAP (SNAP) with TGN38 (n=3-4; 71-91 cells per condition). (**F**) Model depicting metabolic regulation of ER redox homeostasis and proinsulin export via NADPH production and glutathione reduction created by Biorender. (**A-C**, **E**) Data represent the mean ± S.E.M. * p < 0.05 by two-way ANOVA with Sidak post-test analysis (**A**, **E**), one-way ANOVA with Dunnet post-test analysis (**C**), or Student t test (**B**).

Cellular reducing equivalents, such as GSH, are re-generated by NADPH-dependent enzymes. Based on this, we reasoned that NADPH production may be necessary to support ER functions, such as proinsulin disulfide bond formation and export, via supplying cellular (GSH) reducing equivalents (Figure 7A) as previously suggested (Gansemer et al., 2020). To test this, islets were treated with recombinant adenovirus expressing either a non-targeting control shRNA (SAFE) or an shRNA targeting Idh1, the major cytosolic NADPH-producing enzyme in β-cells, which has been previously shown to support insulin granule exocytosis (Bauchle et al., 2021; Ferdaoussi et al., 2015; Ronnebaum et al., 2006). Note that Idh1 loss does not impair mitochondrial function (Ronnebaum et al., 2006). By qRT-PCR, we observed a strong suppression of Idh1 (Figure 7B). Importantly, our viral backbone co-expresses mCherry from a separate RNA pol II promoter (PGK) allowing the specific identification of transduced (Idh1 or SAFE shRNA) islet cells (Bearrows et al., 2019). Using this system, we examined insulin granule formation via fluorescent pulse-chase labeling of our proCpepSNAP reporter, which is under control of the rat insulin promoter (RIP), such that mCherry+ proCpepSNAP+ cells correspond to Idh1 knockdown β-cells (Figure S5). Here, we observed a substantial (> 50%) decrease in the production of nascent (proCpepSNAP-labeled) insulin granules in Idh1 knockdown compared to control (SAFE) β-cells (Figure 7C, D). Next, we examined whether DTT addition could replenish necessary reducing equivalents and thereby restore the deficit in insulin granule formation in Idh1 knockdown β-cells (Figure 7A). Prior to pulse-chase labeling, mouse islets were treated with 0.5 mM DTT (or vehicle control) for 4 h. Consistent with our previous data (Figure 6D, E), addition of DTT, compared to vehicle control, did not affect insulin granule formation in Ad-shRNA-SAFE treated control cells (Figure 7E, F). In contrast, pre-treatment of Idh1 shRNA KD β-cells with DTT rescued insulin granule formation, restoring the levels to the Ad-shRNA control. Collectively, our data demonstrate that proinsulin trafficking and insulin granule formation can be regulated by NADPH production and reducing equivalent availability (Figure 7A) and highlight a direct link between mitochondrial and secretory dysfunction in the decline of β-cell function in T2D.

**Figure 7.**
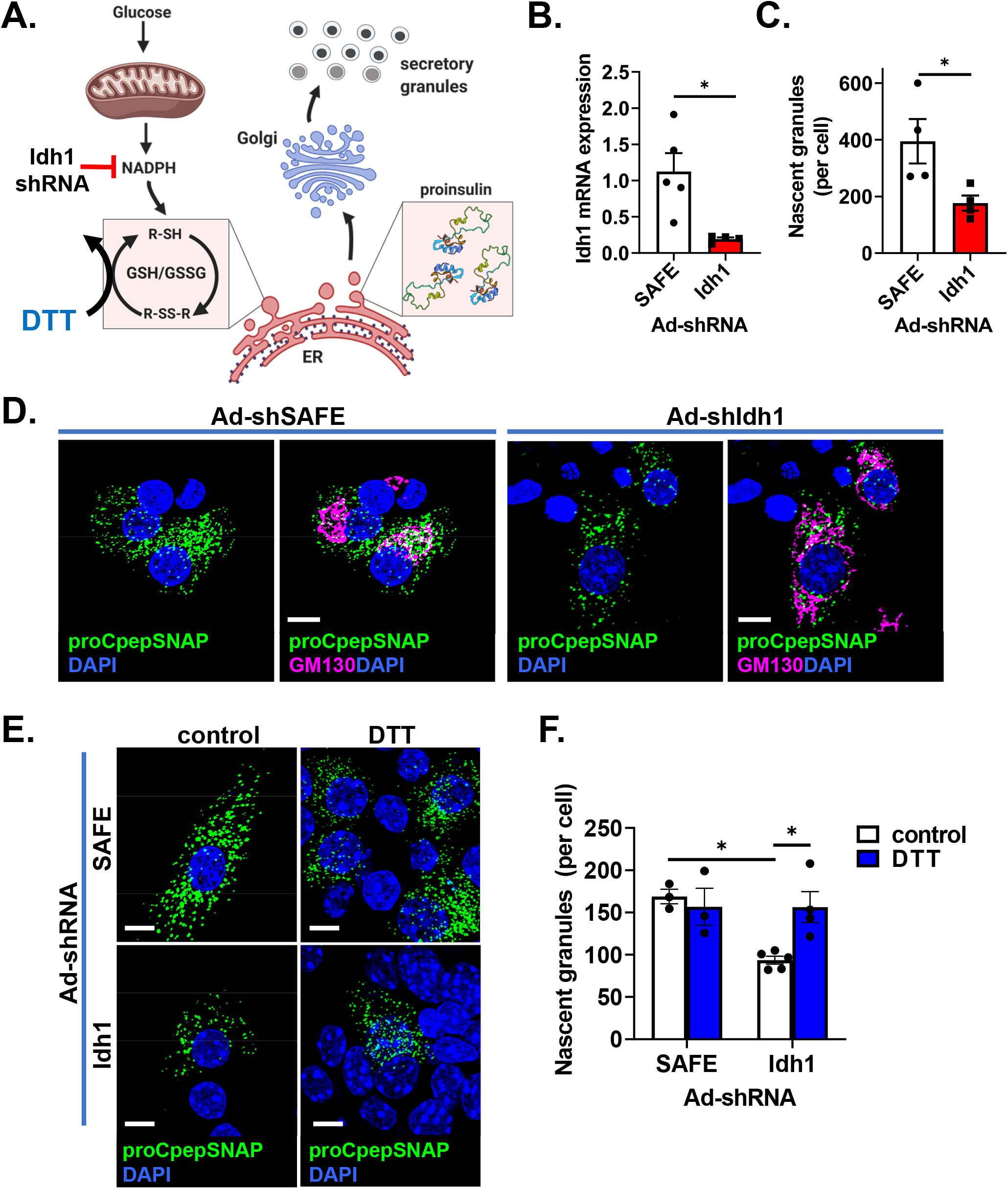
Loss of cytosolic NADPH-producing enzyme Idh1 impairs insulin granule formation. (**A**) Model depicting metabolic regulation of ER redox homeostasis and proinsulin export via NADPH production and glutathione reduction created by Biorender. **(B**-**F**) Mouse islets were co-treated with AdRIP-proCpepSNAP and either Ad-shRNA-SAFE (control) or Ad-shRNA-Idh1 and analyzed after 72 h. (**B**) Idh1 mRNA expression was determined by qRT-PCR (n=4-5). (**C-F**) Mouse islets were pulse-labeled with SNAP-505 for 20 min (green) and chased for 2 h. Cells were fixed, immunostained for GM130 (magenta), and counterstained with DAPI (blue). mCherry expression was used to identify shRNA-expressing cells in each condition. Total number of nascent (proCpepSNAP-labeled) granules per mCherry+ cell was quantified (**C**) (n=4; 4-11 cells per mouse) and representative images shown (**D**) (scale bar = 2 μm). (**E, F**) Mouse islets were treated with vehicle (control) or DTT (0.5 mM) for 4 h prior to being pulse-labeled. Representative images (**E**) are shown from pulse-chase labeling and quantified per mCherry+ cell (**F**) (n=3-5; 32-54 cells per condition). (**B, C, F**) Data represent the mean ± S.E.M. * p < 0.05 by Student t test (**B**, **C**) or two-way ANOVA with Sidak post-test analysis (**F**).

## Discussion

Defects in the β-cell’s secretory pathway, including changes to insulin trafficking, reduced insulin storage, impaired proinsulin processing, and hyperproinsulinemia, have been known for decades to occur in T2D (Alarcon et al., 2016; Halban et al., 2014; Kahn et al., 2009; Kahn and Halban, 1997; Like and Chick, 1970; Masini et al., 2012; Ward et al., 1984). Increased exocytosis of immature granules (Rhodes and Alarcon, 1994) as well as enhanced degradation of newly synthesized insulin granules in T2D (Pasquier et al., 2019; Shrestha et al., 2020) contribute to the overall decrease in insulin storage and hyperproinsulinemia; however, the underlying mechanisms for these defects are poorly understood and the links to overnutrition and hyperglycemia remain vague. Our study focuses on an emerging concept that oxidative protein folding in the ER is perturbed in the pathogenesis of β-cell dysfunction (Arunagiri et al., 2019; Jang et al., 2019; Tran et al., 2020) and we link this defect to the loss of insulin granules. We show that changes in cellular redox homeostasis delay proinsulin export from the ER leading to decreased insulin granule formation in animal and cell culture models of overnutrition and diabetes. Furthermore, we demonstrate that ER export of proinsulin can be regulated by NADPH and reducing equivalent availability. Loss of the NADPH-producing enzyme, Idh1, compromises insulin granule formation, while addition of chemical reducing equivalents can restore insulin granule production. Collectively, our data highlight a direct link between nutrient metabolism and the β-cell’s secretory pathway as a critical node in the regulation of insulin biosynthesis (Figure 8)

**Figure 8.**
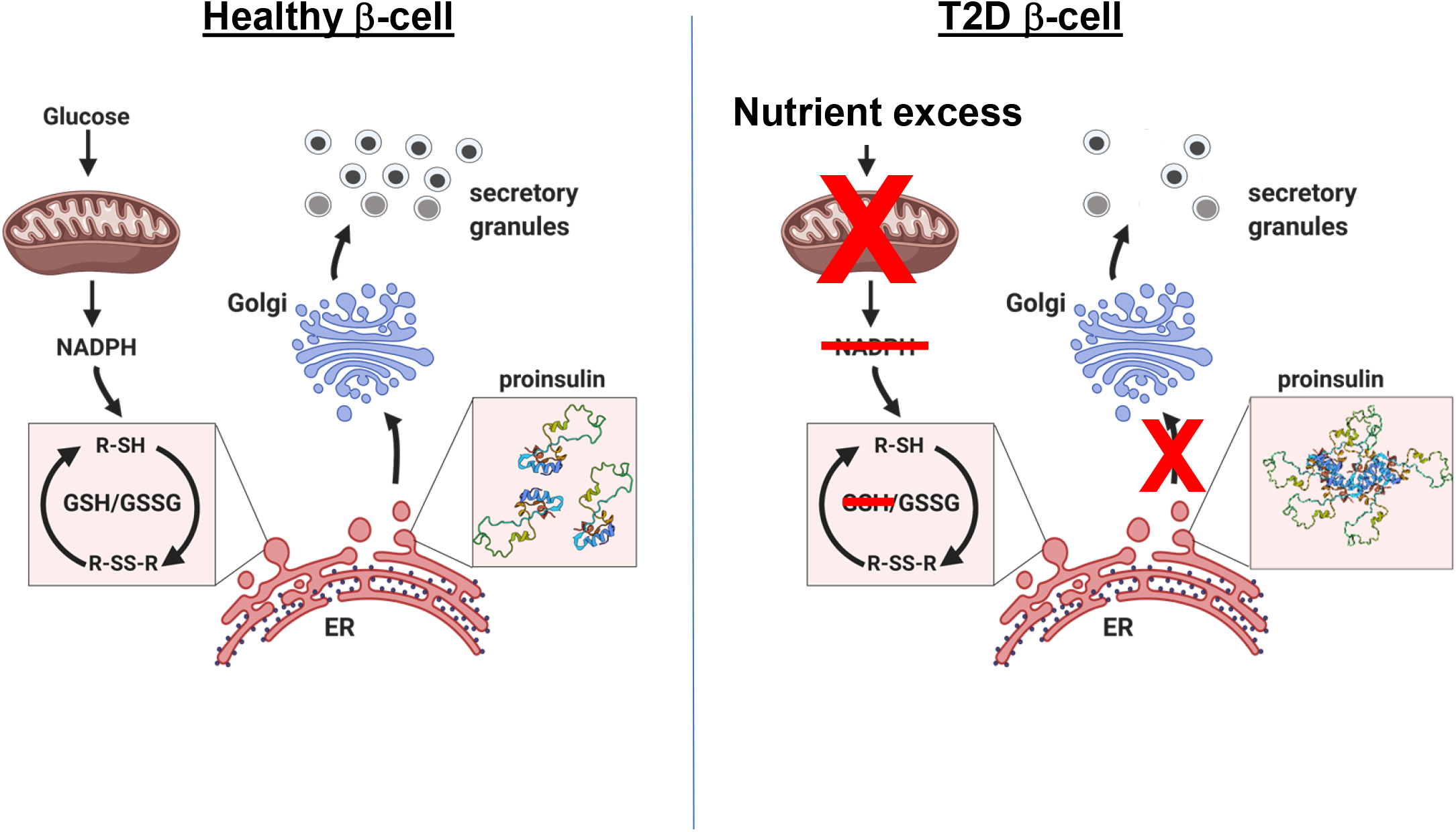
Model of metabolism-coupled proinsulin trafficking. In this model, glucose metabolism by the mitochondria generates NADPH, which is used to regulate glutathione cycling (GSH/GSSG) and produce GSH reducing equivalents. GSH is necessary to maintain sequential rounds of protein disulfide bond isomerization (R-SH, R-SS-R) in the ER to facilitate proinsulin folding. Properly folded proinsulin can be exported from the ER to the Golgi, where it is packaged into immature secretory granules. In response to nutrient excess in T2D, mitochondrial dysfunction leads to a decrease in NADPH and an insufficient supply of GSH reducing equivalents to sustain proinsulin folding. This leads to a delay in proinsulin ER export and ultimately a reduction in the production of insulin secretory granules. Proinsulin NMR structure (2KQP) was derived from the RCSB protein database. This model was created with Biorender.com.

Following the emergence of nascent proteins from the ER protein-conducting channel, Sec61, disulfide bonds readily form in the oxidizing environment of the ER lumen. In the case of disulfides arising from non-sequential cysteine residues, such as the three disulfide bonds in proinsulin, rearrangement of mis-paired disulfides requires reduction via PDIs and subsequent re-oxidation of cysteine thiols to form the correct disulfide linkages (Ellgaard et al., 2018; Winter et al., 2002). Using our proximity labeling system, we identified 11 ER oxidoreductases, including Prdx4 and Pdia1, that interact with proinsulin under normal physiological conditions, highlighting a potential critical role for disulfide bond isomerization in proinsulin folding. Following chronic exposure to hyperglycemic and hyperlipidemic-like conditions, we observed a significant elevation in the interactions between proinsulin and ER oxidoreductases, which has been previously reported in human T2D β-cells (Tran et al., 2020). This increase in proinsulin interactions with ER oxidoreductases may be driven by the aberrant formation of mis-paired intermolecular disulfide bonds between proinsulin molecules (Arunagiri et al., 2019; Tran et al., 2020) and lead to the ER export delay reported here. Based on these data, we propose that the ER lumen becomes hyperoxidizing during the pathogenesis of β-cell dysfunction, which impedes the necessary re-shuttling of proinsulin’s disulfide linkages and thereby delays ER export of proinsulin (Figure 8). In support of this mechanism, the delay in ER export can be reversed by the addition of chemical reducing equivalents, suggesting that proinsulin disulfide bond isomerization is a kinetic determinant in proinsulin export from the ER. Similarly, antioxidant treatment has been shown to decrease ER stress and attenuate β-cell dysfunction and diabetes onset in genetic mouse models of ER dysfunction (Han et al., 2015; Hassler et al., 2015; Malhotra et al., 2008). Interestingly, we observed no defect in ER to Golgi transit of CgB, which contains a single pair of sequential cysteines forming its lone disulfide linkage. Thus, CgB may be less dependent on reducing equivalents for correct disulfide bond formation and ER export. Together, our data provide compelling evidence linking defects in oxidative protein folding in the ER to a decrease in insulin granule formation via delayed ER export of proinsulin during the progression of β-cell dysfunction in T2D (Figure 8).

Recent data demonstrates that ER oxidative protein folding can be regulated via nutrient metabolism (Gansemer et al., 2020) through NAPDH dependent recycling of redox intermediates, glutathione and/or thioredoxin, which are necessary to reset PDIs for sequential rounds of client protein disulfide bond isomerization (Birk et al., 2013; Ellgaard et al., 2018). Distinct from other cell types, the β-cell is uniquely positioned to sense physiological fluctuations in glucose via the generation of metabolic signals (Boland et al., 2017). Here, we speculate that nutrient metabolism in the β-cell is directly coupled to secretory function through the regulation of ER redox poise. In this pathway, a rise in glucose metabolism, which increases preproinsulin mRNA translation (Wicksteed et al., 2003), also supplies reducing equivalents to facilitate proinsulin disulfide bond isomerization in the ER and thereby expedite proinsulin biosynthesis and insulin production. In support of this, loss of glucose-dependent NADPH production via Idh1 suppression impairs insulin granule formation, which can be restored via replacement of cellular reducing equivalents. During the pathogenesis of T2D, chronic exposure to nutrient excess and hyperglycemia has the unfortunate consequence of impairing mitochondrial function^45^ resulting in diminished production of NADPH and glutathione/ thioredoxin reducing equivalents (Ferdaoussi et al., 2015), which are necessary for ER function (Birk et al., 2013; Ellgaard et al., 2018). As a result, the ER lumenal environment falters in efficient proinsulin disulfide bond isomerization leading to the formation of proinsulin aggregates (Arunagiri et al., 2019; Tran et al., 2020), which may trigger ERAD and other degradative pathways (Pasquier et al., 2019; Shrestha et al., 2020, p. 1), and ultimately decrease insulin granule formation. While we failed to detect increased levels of cellular ROS, we cannot rule out the possibility that a portion of the reduced glutathione pool, was consumed to buffer accumulating ROS and thereby lead to a decrease in GSH available for proinsulin disulfide bond isomerization. Similarly, the limited supply of NADPH could stem directly from decreased mitochondrial output of necessary substrates such as citrate/isocitrate and/or increased utilization by NADPH oxidases to elevate cytosolic (and possibly ER) levels of hydrogen peroxide. Based on these possibilities, future studies examining the mechanisms intertwining β-cell nutrient metabolism and secretory function may offer unique insight into novel pathways that could be exploited for diabetes treatment.

Numerous studies have demonstrated that induction of ER stress can perturb β-cell function and may contribute to the development of β-cell dysfunction in T2D (Yong et al., 2021). Loss of critical ER stress response sensors, such as PERK and Ire1α, can sensitize β-cells to nutrient stresses and/or directly impair β-cell function leading to the development of diabetes (Delépine et al., 2000, p. 2; Harding et al., 2001; Hassler et al., 2015; Xu et al., 2014). Similarly, loss of the ER chaperone, GRP94, the BiP co-chaperone, p58^IPK^, and PDIs, Pdia1 and Prdx4, compromise proinsulin folding, resulting in ER stress and loss of insulin content (Ghiasi et al., 2019; Jang et al., 2019; Kim et al., 2018, p. 94). In humans, monogenic forms of diabetes arising from *INS* mutations can prevent proper proinsulin disulfide bond formation acting as dominant negative suppressors of wild type proinsulin trafficking (Haataja et al., 2021; Liu et al., 2007; Støy et al., 2010). While these studies demonstrate that perturbations in ER functions and proinsulin folding can compromise β-cell health and insulin production, direct evidence for ER stress as a causative feature in the development of T2D is less clear (Alarcon et al., 2016; Marchetti et al., 2007). Increased PERK and Ire1α activity have been described in pre-diabetic β-cells (Engin et al., 2014; Lipson et al., 2006; Yong et al., 2021), yet diminished, rather than increased, expression of ATF6 and XBP-1(s) occur in β-cells from humans with long-standing T2D (Engin et al., 2014). Moreover, human β-cells are known to cycle between varying levels of UPR protein expression (Xin et al., 2018), which may represent the natural β-cell adaptation to changes in nutrient load. Indeed, activation of the ER stress pathway is necessary for the β-cell proliferative response to diet-induced obesity (Sharma et al., 2015). Expansion of ER membranes has been demonstrated in both human and rodent T2D β-cells, which may be necessary to support increased proinsulin biosynthesis (Alarcon et al., 2016; Boland et al., 2017; Marchetti et al., 2007; Masini et al., 2012); however distension and/or dilation of the ER lumen, a classic marker of proteotoxic ER stress, is not commonly observed. Consistent with these data, we failed to detect changes in ER ultrastructure in diabetic db/db mice and saw little evidence for overt ER stress response activation in our cell culture model, despite clear effects on ER retention of proinsulin in both models. These unresolved issues encourage continued investigation in defining how physiological stresses manifest in the β-cell and determining when the adaptive responses become insufficient to mitigate cellular damage and/or become maladaptive.

## Materials and methods

### Cell culture, islet isolation, and reagents

832/3 cells (a kind gift from Dr. Christopher Newgard) were cultured as previously described (Hohmeier et al., 2000). 832/3 cells stably expressing proCpepSNAP have been described previously (Bearrows et al., 2019). For introduction of adenoviral reporters, cells were transduced with ∼1-5 x 10^7^ IFU/ mL adenovirus for 18 h and assayed 72-96 h post-treatment. Cell culture reagents were from Thermo Life Technologies unless specified otherwise. Chemical reagents were from Sigma-Aldrich unless specified otherwise. BSA-conjugated fatty acid solution was prepared as follows: Oleate/palmitate (2:1; 10 mM) was dissolved in water at 95⁰C, cooled to 50⁰C, BSA (Sigma, fatty acid free) added to a final concentration of 10%, and maintained at 37⁰C for an additional 1 h. Mouse islets were isolated via collagenase V digestion and purified using Histopaque 1077 and 1119. Islets were cultured in RPMI supplemented with 10% fetal bovine serum and 1 % penicillin and streptomycin and maintained at 37°C in 5% CO_2_. Pools of islets were transduced with ∼1-5 x 10^8^ IFU/ mL adenovirus for 18 h and assayed 72-96 h post-treatment.

### Animal studies

Male C57BL6/J mice (8-10 wk old; Jackson Laboratories) were placed on a Western diet (Research Diets, D12079B) or standard chow for up to 14 weeks. Body weight and *ad lib* fed blood glucose were measured weekly. BKS.Cg-Dock7^m^ (C57BLKS/J) Lepr^db/+^ and Lepr^db/db^ mice (db/+, db/db) were either generated by heterozygous cross and genotypes confirmed via real time PCR according to Jackson Laboratories or directly purchased from Jackson Laboratories. Hyperglycemic db/db mice (*ad* lib fed blood glucose > 220 mg/dl) were compared to age-matched (10-14 weeks old), normoglycemic littermate controls (db/+). Glucose tolerance was measured in 4-6 hr fasted mice given a 1 mg/ g body weight glucose (i.p.) challenge. Blood glucose was determined using a One Touch Ultra 2 glucometer. Plasma insulin was determined by ELISA (ALPCO). All animal protocols were approved by the University of Iowa Institutional Animal Use and Care Committee

### Plasmids and viruses

The shuttle plasmid, pENTR-RIP, containing a rat insulin promoter (RIP), multiple cloning site, chloramphenicol resistance marker, ccdB, and a bovine growth hormone (bGH) polyA signal was generated by Gibson assembly (IDT) and subcloned into pENTR2b (Thermo Life) replacing the entire cassette between the attP1 and attP2 sites. V5-tagged APEX2 was inserted near the ApaI site of human preproinsulin via gblock synthesis (IDT) and subcloned into pENTR-RIP via Gibson assembly. PCR fragments of CgB (DNASU) and CLIP (gift from Eric Campeau; RRID:Addgene_29650) were subcloned into pENTR-RIP via Gibson assembly. RIP-proCpep-APEX2 and RIP-CgB-CLIP were assembled into a modified pAd-PL/DEST via Gateway cloning using LR Clonase II. Adenovirus containing rat insulin promoter (RIP) expressing Grx1-GFP2 was a kind gift from A. Linneman, Indiana University School of Medicine (Reissaus et al., 2019). Adenoviruses expressing shRNA targeting mouse Idh1 (AdU6-Idh1 shRNA-PGK-mCherry) and non-targeting control shRNA (AdU6-SAFE-shRNA-PGK-mCherry) have been described previously (Bauchle et al., 2021). Recombinant adenoviruses were generated in HEK293 cells and purified by cesium chloride gradient. All sequences were verified by the Iowa Institute of Human Genetics, University of Iowa.

### Glucose-stimulated insulin secretion

Insulin secretion was measured by static incubation as previously described (Stephens et al., 2017). Cells were lysed in RIPA buffer and total protein determined by BCA (Pierce). Insulin (secreted and content) was measured by ELISA (rodent 80-INSMR-CH10; ALPCO) or Alphalisa (AL204C; Perkin-Elmer Insulin).

### Immunoblot analysis

Clarified cell lysates were resolved on 4-12% NuPAGE gels (Thermo Life Technologies) and transferred to supported nitrocellulose membranes (BioRad). Membranes were probed with diluted antibodies raised against chromogranin B (goat, Santa Cruz C-19 1:1000), CHOP (mouse, 1:1000 Cell Signaling 2895), cleaved Caspase 3 (rabbit, R&D systems MAB835, 1:500), CPE (rabbit, Proteintech 13710-1-AP, 1:1000), GRP94 (rabbit, kind gift of Dr. Christopher Nicchitta, Duke University, 1:5000), PC2 (rabbit, Proteintech 10552-1-AP, 1:1000), proinsulin (mouse, Iowa Developmental Studies Hybridoma Bank GS-98A, 1:500), syntaxin-4 (rabbit, Millipore AB5330, 1:1000), TMED9 (rabbit, Proteintech 21620-1-AP, 1:10,000), TMED10 (rabbit, Bethyl Laboratories, A305-219A, 1:1000), gamma-tubulin (mouse, Sigma T5326, 1:10,000). Donkey anti-mouse, anti-rabbit, or anti-goat antibodies (Licor) coupled to IR-dye 680 or 800 were used to detect primary antibodies. Streptavidin-conjugated to IR dye 680 (1:1000, Thermo) was used to detect biotinylated proteins. Blots were developed using an Odyssey CLx Licor Instrument.

### Fluorescence microscopy and imaging

Isolated islets expressing proCpepSNAP (AdRIP) were dispersed into monolayers using Accutase (Sigma-Aldrich) and plated onto HTB9 coated coverslips or 6 cm glass bottom dishes (Mattek) as previously described (Bearrows et al., 2019; Hayes et al., 2017). 832/3 cells stably expressing proCpepSNAP were plated on HTB9 coated coverslips at low density and cultured overnight as previously described (Bearrows et al., 2019). For SNAPtag labeling, cells were initially incubated with SNAPcell block (10 µM; NEB) diluted in culture media for 20 min, washed 3 times for 5 min each. To allow for *de novo* protein synthesis of new proCpepSNAP, cells were cultured for an additional 2 h in experiments examining post-Golgi granule formation or 45 min in experiments examining ER-Golgi transport. Cells were pulse-labeled with SNAPcell-505 (1 µM; NEB) for 20 min in media, washed 3 times for 5 min each in culture media with glucose (5 mM) and chased as indicated. For studies utilizing CgB-CLIP, an identical labeling scheme was followed using CLIPcell block (10 µM; NEB) and CLIPcell-TMR (1 µM; NEB) where appropriate. Following treatments, cells were fixed in 10 % neutral-buffered formalin. For immunostaining, cells were incubated overnight with antibodies raised against GM130 (mouse, BD Transduction 610823, 1:200), GRP94 (rabbit, kind gift of Dr. Christopher Nicchita, Duke University, 1:500), TGN38 (mouse, Novus Biologicals NB300-575, 1:200), or V5 epitope tag (rabbit, GeneTex GTX117997, 1:100) as indicated. Highly cross-adsorbed fluorescent conjugated secondary antibodies (whole IgG, donkey anti-guinea pig-AlexaFluor 488, donkey anti-rabbit rhodamine red-X, donkey anti-mouse AlexaFluor 647; Jackson ImmunoResearch) were used for detection. Cells were counterstained with DAPI and mounted using Fluorosave (Calbiochem). Images were captured on a Leica SP8 confocal microscope using a HC PL APO CS2 63x/1.40 oil objective with 3x zoom as z-stacks (5 per set, 0.3 μm step, 0.88 μm optical section) and deconvolved (Huygen’s Professional). Granule distance measurements from the Golgi were determined using a distance transformation module in Imaris (Bitplane) from spot-rendered granules (SNAP-505 labeled) and surface rendering of the Golgi identified by GM130 or TGN38 immunostaining. Granule distances were binned and expressed as a percentage of the total to normalize between cells. Total granule number were also calculated from these studies with respect to the number of pulse-chased labeled cells. For islet studies, we used a cutoff of less than 0.5 μm to identify granules proximal to the Golgi as attached, and granules > 2 μm as successful Golgi export based on time course studies of granule distance clustering (Bearrows et al., 2019). Golgi/ER fluorescence of SNAP labeling was determined from thresholded masks defining Golgi and ER area by immunostaining using macros written for Fiji NIH software. For plasma membrane detection, immunostained (fixed) cells were maintained in PBS and imaged using a Leica TIRF AM microscope (100x oil objective) in TIRF mode with a penetration depth of 110-150 nm. In some experiments, SNAP25 immunostaining was used to define the plasma membrane. Granule numbers from SNAP-labeling, cell area and or cell number (defined by nuclei) were determined by Fiji/ImageJ.

For glutathione (GSSG/GSH) redox measurements, 832/3 cells were treated with AdRIP-Grx1-GFP2 and imaged on an inverted Olympus IX83 microscope with a 20x objective (HC PL APO CS2; 0.75 NA) using Chroma 49002 GFP/CY2 Bandpass (470/535) and custom Chroma BX3-mounted (395/510) filters. Cells were subsequently cultured and re-imaged following DTT (10 mM) and diamide (5 mM) treatment, for 12 min each (Bauchle et al., 2021). Fluorescent intensities of each channel were calculated from masked images (to remove background) of whole cells using macros written for Fiji NIH software. Ratiometric intensities (395 nm/470 nm) were normalized by comparing to DTT (0%) and diamide (100%) treated samples using GraphPad Prism software.

For ROS measurements, 832/3 cells were cultured in opaque HTB9-coated 96-well plates as indicated. Menadione (100 μM) treatment for 1 h was used as a positive control. Cells were incubated for 0.5 h with Cell ROX Deep Red (5 μM), washed, and analyzed using a Spectromax MD5 plate reader.

### Ultrastructure

All EM related reagents were from Electron Microscopy Sciences (EMS; Hatfield, PA). Isolated islets were PBS washed and fixed in 2.5% glutaraldehyde, 4 % formaldehyde cacodylate buffer overnight (16-18 h) at 4⁰C. Tissue was post-fixed in 1% OsO_4_ for 1 h, dehydrated using a graded alcohol series followed by propylene oxide and embedded in Epon resin as previously described (Joshi et al., 2014). Resin blocks were cut to ultrathin (50-70 nm) sections with a diamond knife and mounted on Formvar-coated copper grids. Grids were double contrasted with 2% uranyl acetate then with lead citrate. Images were captured at 1,500x, 3,000x, 6,000x, and 8,000x magnifications by a JEOL JEM-1400 transmission electron microscope.

### Density gradient isolation of secretory granules

Cells were collected, washed in ice-cold PBS, and disrupted using 15 strokes in a pre-chilled ball-bearing cell homogenizer (Isobiotec) with a 14 µm clearance in 10 mM MES pH 6.5, 1 Mm MgSO4, 1 mM EDTA, 0.3 M sucrose as previously described (Bearrows et al., 2019; Stephens et al., 2017). Post-nuclear supernatants were layered atop 8-23% linear iodixanol (Optiprep) gradients and resolved at 110,000 x g’s in an SW41 for 16-18 h. Fractions were manually collected by tube puncture. Iodixanol gradients were verified by absorbance at 340 nm. Insulin content was determined by ELISA (ALPCO) and normalized to the total insulin content from post-nuclear supernatant.

### APEX2 biotinylation and proteomics analysis

APEX2 biotinylation was performed using a modified protocol (Hung et al., 2016) as follows. Cells expressing proCpepAPEX2 were cultured for 1 h in SAB containing 2.5 mM Glc followed by addition of 500 µM biotin phenol (Adipogen) for 0.5 h. Diluted H_2_O_2_ was added to a final concentration of 1 mM and incubated for 1 min. Non-peroxide-treated samples were used as controls for non-enzymatic labeling such as endogenous biotin-containing proteins. Cells were washed 5x in PBS containing 10 mM sodium ascorbate, 5 mM Trolox (Acros), and 10 mM sodium azide for 1 min each. Cells were scraped, collected by centrifugation, and lysed in RIPA containing 10 mM sodium ascorbate, 5 mM Trolox, 10 mM sodium azide, and Halt protease cocktail (Thermo). For proteomics studies, cell pellets were snap frozen in a dry-ice ethanol bath prior to lysis. Pierce-660 (Thermo Life) was used to determine protein concentrations. For pulldowns, clarified cell lysates (1-2 mg/mL) were added to high capacity Neutravidin agarose resin (Thermo Life) and rotated end over end overnight (16-18 h) at 4⁰C. Bound proteins were washed in RIPA, 1 M KCl, 0.1 M sodium carbonate, 2 M urea in 10 mM Tris pH 8.0, RIPA solution without detergent, and eluted at 70⁰C for 15 min in 2x LDS sample buffer (Thermo Life) containing 2 mM biotin.

Proteomic analysis was performed on Neutravidin affinity-purified lysates (∼1.5 mg starting material) with reproducible recovery verified with 4-12% NuPAGE (Thermo Life) and silver staining. Sample pre-cleaning and in-solution tryptic digestions were accomplished with a lab-built silica / C-18 trapping mini-column (Zougman et al., 2014). Recovered peptides were labeled by reductive amination as previously described (Yu et al., 2017) using distinct (light, medium, heavy) dimethyl isotopomers (Boersema et al., 2009; Hsu et al., 2003; Yu et al., 2017). Completion of labeling was confirmed by MALDI-TOF and distinct isotopomer-labeled samples from each treatment were mixed at equal ratios for LC–MS/MS analysis. Rather than subtracting the median ratio of spatially inaccessible false positives to normalize regional APEX, heavy (+36) and light (+28) isotopes were applied in crossover to two replicates of samples cultured in OPG vs BSA control. Non-peroxide treated (no APEX labeling) cells were labeled with the medium isotopic channel (+32). Mass spectrometry data were collected using an QExactive hf mass spectrometer coupled to an Easy-nLC-1200™ system. The autosampler typically loads 0.3 µg of reconstituted digest on a 2.5cm C18 trap (New Objective, P/N IT100-25H002) coupled to waste or a 200cm x 75um pillar array column from PharmaFluidics. The column outlet floated at high voltage via a gold connector secured to the micro-cross assembly (IDEX, P/N UH-752). MS1 scans were acquired every 3 sec. with automatic gain control set to 3e05 and 60,000 resolution. Ions with 2 to 6 charges were isolated within a 1.2 Th pass window and targeted for HCD activation. The AGC target for HCD was 4E05, 30% NCE, using 75 ms maximum fill time. and 30k resolution at 400 m/z. Searches were performed with both MaxQuant, version 1.6.11 and Byonic search engines (Protein Metrics ver. 2.8.2). Search databases were composed of the Uniprot KB for rat (June 11, 2019). Searches were set to accept static ReDi labels and +57 on Cys residues. Variable oxidation was assumed to occur at Met only. Quantification was achieved by comparing the integrated intensity of each MS1 channel in a sequence-matched triplet derived from the multiplexed sample. Scaffold was used to collate fold enrichment of BSA- and oleate/palmitate + glucose-cultured samples compared to non-peroxide-treated samples. Classifiers to assign active engagement in ISG assembly were established by ROC curve analysis as described(Hung et al., 2016). The log_2_ fold enrichment in the peroxide-treated (BSA) samples relative to no peroxide was determined and was generally 2.3 or more for biotinylated proteins. For proteins with a log_2_ fold change of 1.2 or greater enrichment in the BSA (control) cultured samples, the relative fold change to oleate/palmitate + glucose cultured samples was determined. Protein subcellular locations were determined using annotations from the rat and human orthologs in the UniProt database and additional confirmation by literature searches as needed.

### Metabolic studies

Mitochondrial function was evaluated using Agilent Seahorse XF96 Extracellular Flux Analyzer. 832/3 cells were seeded at a density of 40,000 cells/well in XF 96-well microplates. Cells were cultured for 72 h in BSA (control) or OPG media as indicated. Prior to the assay, cells were washed and incubated in serum-free DMEM supplemented with 5mM HEPES, 2 mM L-glutamine and 1 mM Na pyruvate (Life Technologies), pH 7.40 with either 2 mM or 20 mM glucose. Mitochondrial stress tests were initiated then cellular respiration was evaluated by monitoring the oxygen consumption rate (OCR). A basal rate was established, followed by addition of 2.5 µM oligomycin A, 1.25 µM FCCP, and 2 µM rotenone and antimycin, each as sequential injections. Data were normalized following post-experiment cell enumeration.

### Quantitative RT-PCR

RNA was harvested using a Zymo RNA minikit and cDNA synthesized in an iScript reaction (Bio Rad). Real-time PCRs were performed using the ABI 7700 sequence detection system and software (Applied Biosystems). All primer sequences are available upon request.

### Statistical Analysis

Data are presented as the mean ± S.E.M. For statistical significance determinations, data were analyzed by the two-tailed unpaired, Student’s t test or by ANOVA with post-hoc analysis for multiple group comparisons as indicated (GraphPad Prism). A *p*-value < 0.05 was considered significant.

## Author contributions

K.E.R., C.K.B., S.C.B. M.R.M., W.S.E., and S.B.S conceived and designed studies. K.E.R., C.K.B., S.C.B. M.R.M., W.S.E., C.J.B., S.E.B. and S.B.S performed the experiments and analyzed the data. C.Y. and M.R.P. performed proteomics studies. K.E.R., S.C.B. and S.B.S. analyzed proteomics data. J.Z. and Y.W. performed TEM. K.E.R. and S.B.S. wrote the manuscript.

## Acknowledgements

We would like to thank Dr. Thomas Rutkowski, Erica Gansemer, and Dr. Ling Yang for helpful comments and discussion and McKenzie Becker for expert technical assistance. We would like to acknowledge the use of the University of Iowa Central Microscopy Research Facility, University of Iowa Proteomics Facility, and University of Iowa Radiation and Free Radical Research Core Facility. This work was supported by startup funds provided by the Fraternal Order of Eagles Diabetes Research Center, University of Iowa to S.B.S., Department of Defense CDMRP grant PR190353 to S.B.S., National Institutes of Health grant R35GM130331 to Y.W., and a National Institutes of Health Predoctoral Training Grant T32GM008629, PI Daniel Eberl to K.E.R. University of Iowa Electron Spin Resonance (ESR) Facility, Holden Comprehensive Cancer Center, is supported by NCI P30 CA086862.

## Competing Interests

No competing interests declared.

## Abbreviations

Biotin phenol (BP); dithiothreitol (DTT); glucose (Glc); glucose-stimulated insulin secretion (GSIS); infectious units (IFU); oleate/palmitate (OP); oleate/palmitate + Glc (OPG); oxygen consumption rate (OCR); protein disulfide isomerase (PDI); rat insulin promoter (RIP); Type 2 diabetes (T2D); untranslated region (UTR)

## Data Availability

The datasets generated during and/or analyzed during the current study are available from the corresponding author upon reasonable request. The mass spectrometry proteomics data have been deposited to the ProteomeXchange Consortium via the PRIDE(Perez-Riverol et al., 2019) partner repository with the dataset identifier PXD028606.

---

**Figure S1.**
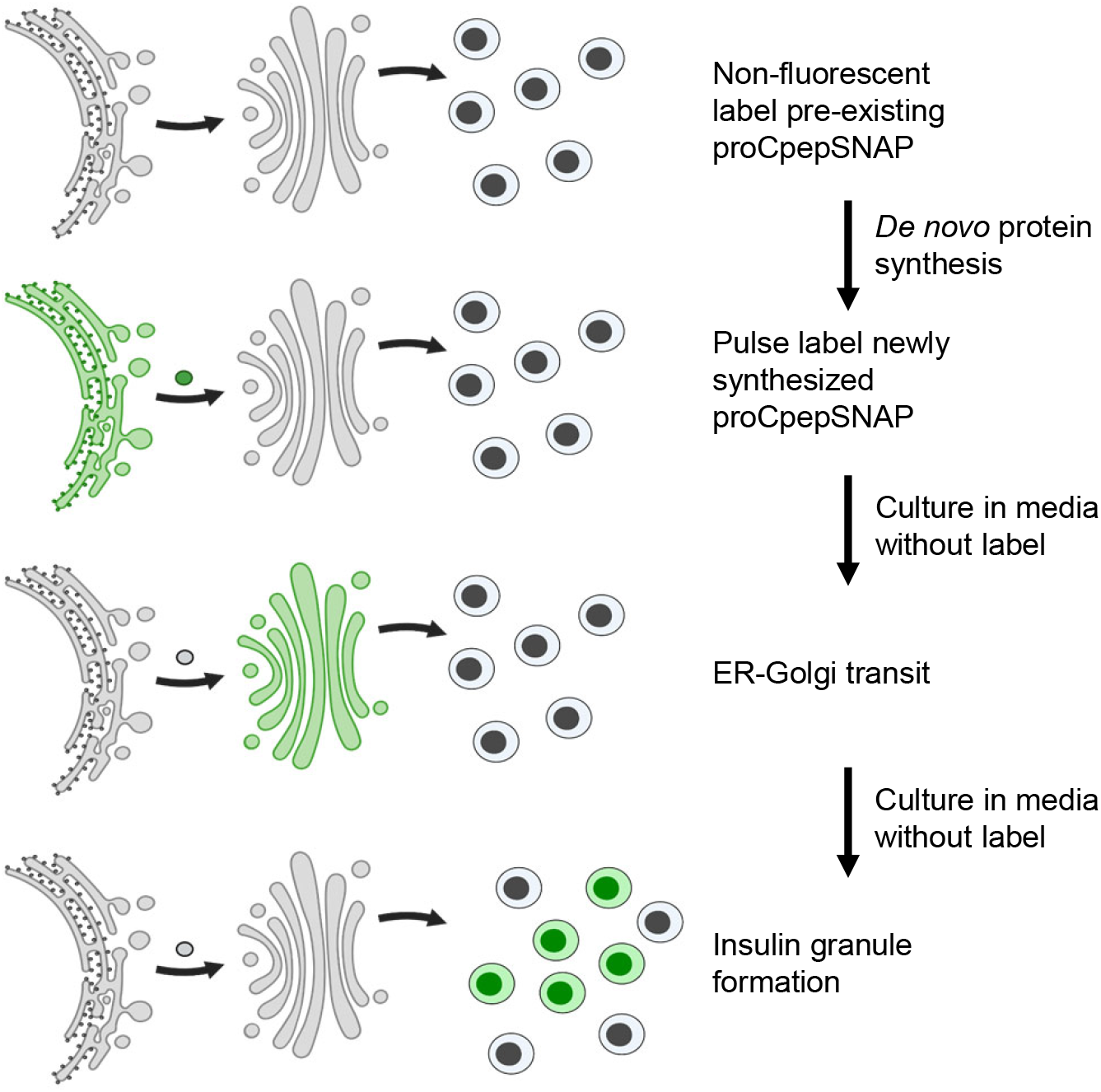
Schematic of proCpepSNAP pulse-chase labeling. To examine proinsulin trafficking, islet cells or 832/3 cells expressing proCpepSNAP are initially treated with SNAPcell block to non-fluorescently label pre-existing proCpepSNAP. Following culture to allow for *de novo* proCpepSNAP synthesis, cells are fluorescently labeled with SNAPcell-505 (green), and then chased in culture media as indicated prior to fixation to examine ER-Golgi transit and post-Golgi insulin granule formation. An analogous labeling scheme is performed for CgB-CLIP. This model was created in Biorender.

**Figure S2.**
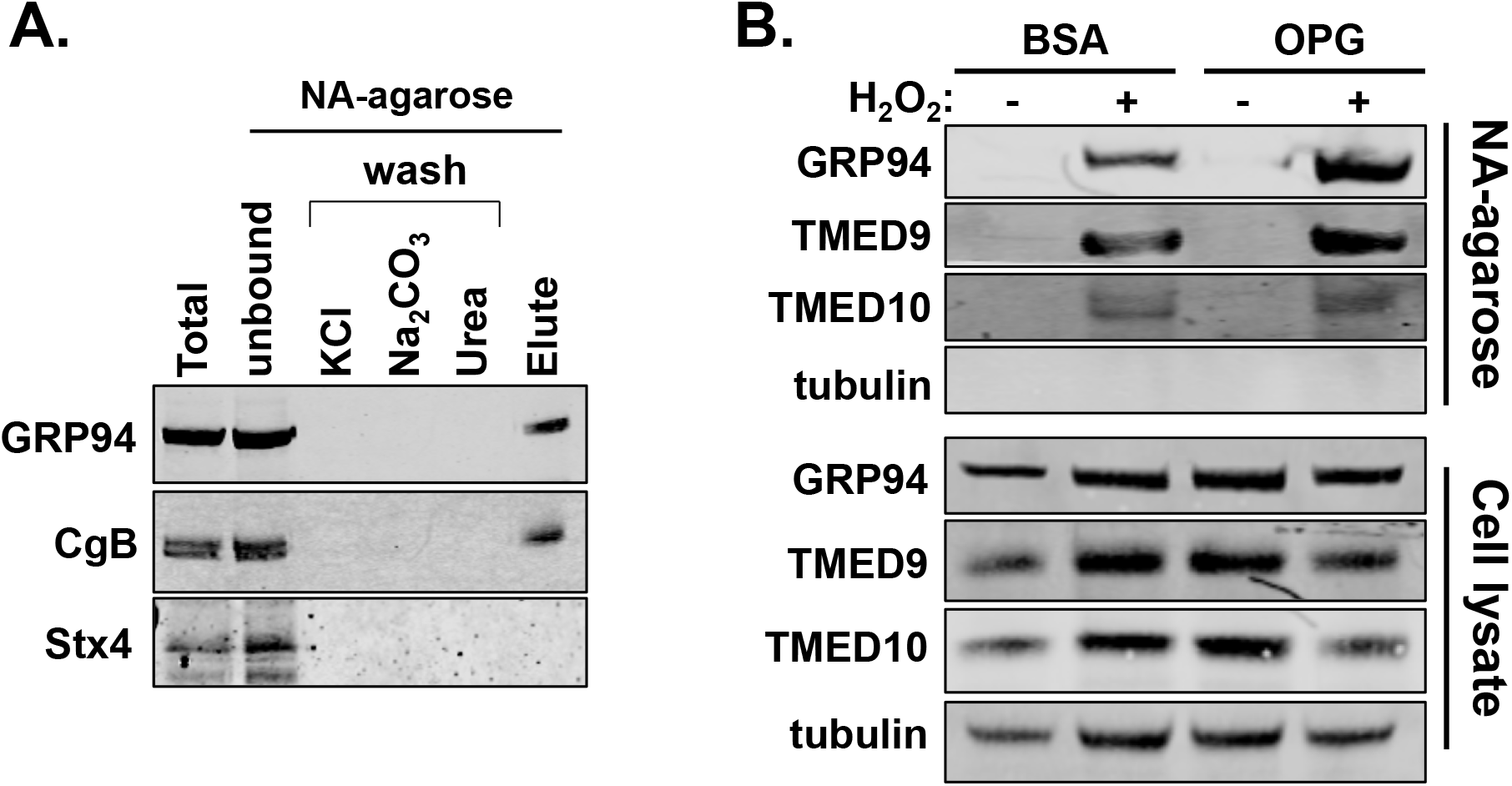
Affinity purification of APEX2 labeling in 832/3 cells. 832/3 cells expressing proCpepAPEX2 were treated with H_2_O_2_ as indicated. (**A**) Lysates from proCpepAPEX2-biotinylated cells were enriched using Neutravidin agarose resin and various stages of purification analyzed by immunoblot probed with GRP94, CgB, and Syntaxin-4 antibodies. (**B**) 832/3 cells expressing proCpepAPEX2 were cultured for 72 h in control media supplemented with BSA or media containing oleate/palmitate (2:1, 1 mM) and elevated glucose (20 mM; OPG) as indicated. Neutravidin agarose-enriched lysates compared to whole cell lysates were analyzed by immunoblot as indicated.

**Figure S3.**
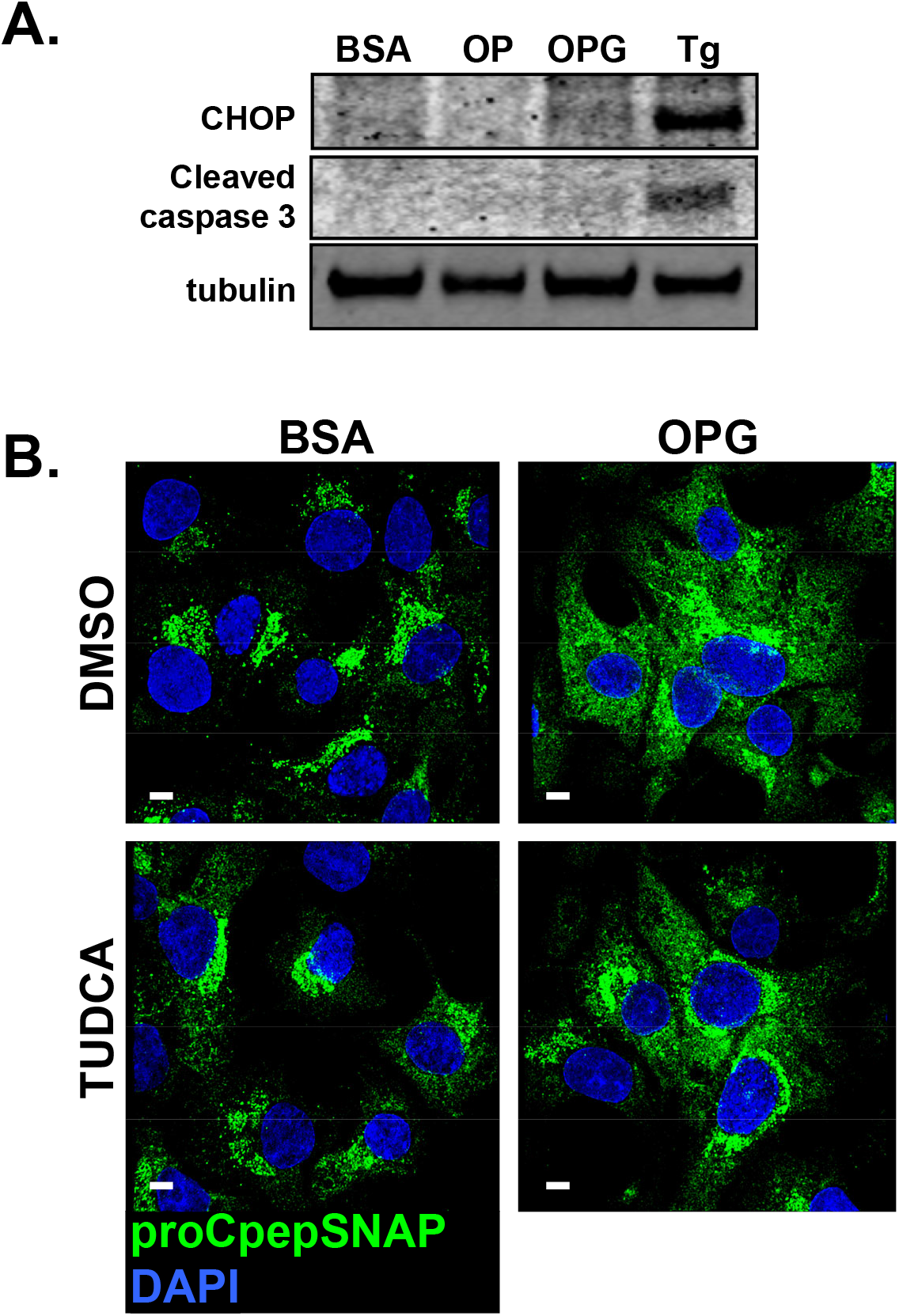
ER export delay of proinsulin is independent of ER stress in OPG-cultured 832/3 cells. 832/3 cells were cultured for 72 h in control media supplemented with BSA, media containing oleate:palmitate (2:1, 1 mM OP) or media containing oleate:palmitate (2:1, 1 mM) and elevated glucose (20 mM; OPG) as indicated. Cell lysates were analyzed by immunoblot compared to cells treated for 18 h with thapsigargin (50 nM) as indicated. (**B**) 832/3 cells stably expressing proCpepSNAP were pretreated with TUDCA (10 μM) or vehicle control (DMSO) for 4 h, as indicated, prior to being pulse-labeled with SNAP-505 for 20 min (green), fixed, and counterstained with DAPI (blue). Representative images are shown (scale bar = 5 μm).

**Figure S4.**
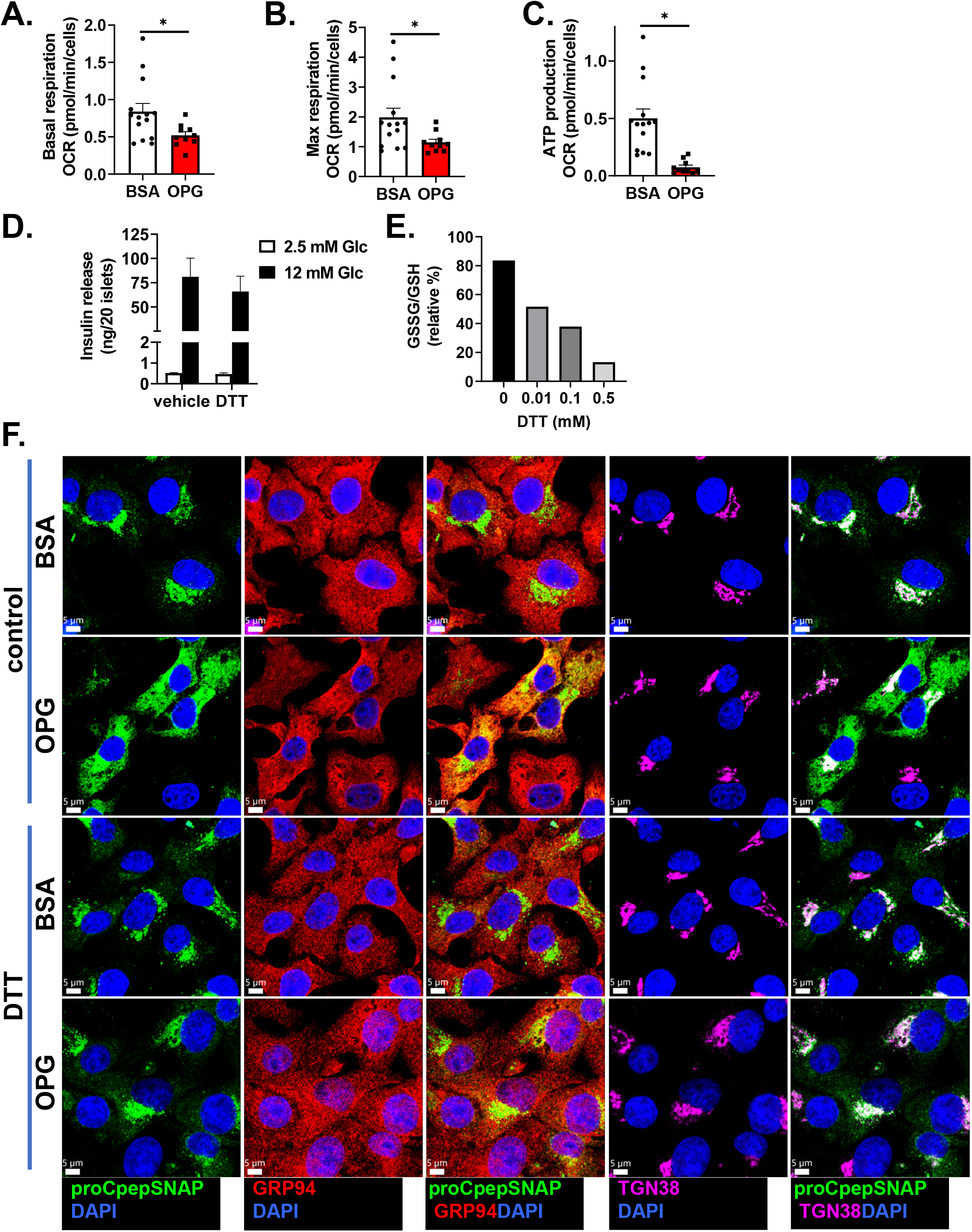
DTT treatment restores ER proinsulin export delay in OPG-cultured 832/3 cells. (**A**-**C**, **F**) 832/3 cells were cultured for 72 h in control media supplemented with BSA (1%) or media containing oleate/palmitate (2:1, 1 mM) and elevated glucose (20 mM; OPG) as indicated. (**A**-**C**) Oxygen consumption rate was measured at stimulatory glucose (20 mM) and following sequential addition of oligomycin (2.5 µM), FCCP (1.25 µM), and rotenone plus antimycin A (2 µM). Basal respiration (**A**), maximum respiration (**B**), and ATP production (**C**) were calculated (n=3, up to 15 replicates per condition). (**D**) Isolated islets (n=4 mice) were cultured overnight with vehicle or DTT (0.5 mM). Glucose-stimulated insulin secretion was measured by static incubation in media containing 2.5 mM Glc followed by 12 mM Glc for 1 h each. (**E**) 832/3 cells expressing Grx1-roGFP2 were treated with the indicated concentrations of DTT for 12 min and imaged at 470/535 nm and 395/510 nm. Ratiometric intensities were normalized to cells treated with DTT (1mM; 0%) and diamide (5 μM; 100%). (**F**) 832/3 cells stably expressing proCpepSNAP were treated with DTT (0.5 mM) for 4 h prior to being pulse-labeled with SNAP-505 for 20 min (green), fixed, and immunostained for TGN38 (magenta), and counterstained with DAPI (blue). Scale bar = 5 μm. Full panels shown of images provided in Figure 6D. (**A**-**D**) Data represent the mean ± S.E.M. * p < 0.05 by Student t test (**A**-**C**).

**Figure S5.**
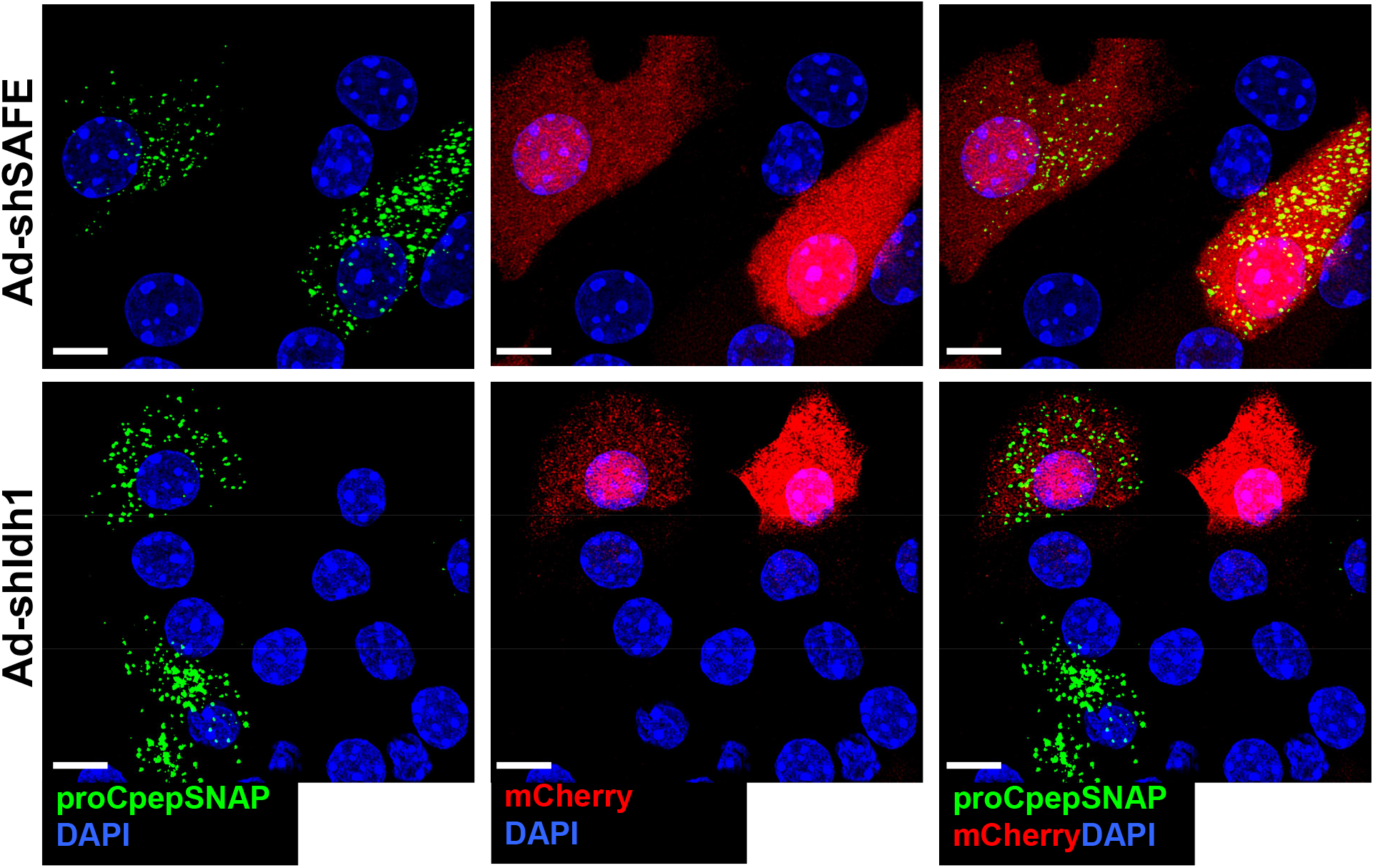
Dual mCherry+ proCpepSNAP+ reporter expression in primary mouse β-cells. Mouse islets were co-treated with AdRIP-proCpepSNAP and either Ad-shRNA-SAFE (control) or Ad-shRNA-Idh1 and analyzed after 72 h. Mouse islets were pulse-labeled with SNAP-505 for 20 min (green) and chased for 2 h. Cells were fixed, immunostained for GM130 (magenta), and counterstained with DAPI (blue). mCherry expression was used to identify shRNA-expressing cells in each condition for analysis (scale bar = 2 μm).

## Notes

### Competing Interest Statement

The authors have declared no competing interest.

### Summary of Updates

updated Title updated figures and text

## Literature Cited

1. Alarcon C, Boland BB, Uchizono Y, Moore PC, Peterson B, Rajan S, Rhodes OS, Noske AB, Haataja L, Arvan P, Marsh BJ, Austin J, Rhodes CJ. 2016. Pancreatic β-Cell Adaptive Plasticity in Obesity Increases Insulin Production but Adversely Affects Secretory Function. Diabetes 65:438–450. doi:10.2337/db15-0792

2. Arunagiri A, Haataja L, Cunningham CN, Shrestha N, Tsai B, Qi L, Liu M, Arvan P. 2018. Misfolded proinsulin in the endoplasmic reticulum during development of beta cell failure in diabetes. Ann N Y Acad Sci 1418:5–19. doi:10.1111/nyas.13531

3. Arunagiri A, Haataja L, Pottekat A, Pamenan F, Kim S, Zeltser LM, Paton AW, Paton JC, Tsai B, Itkin-Ansari P, Kaufman RJ, Liu M, Arvan P. 2019. Proinsulin misfolding is an early event in the progression to type 2 diabetes. Elife 8:e44532. doi:10.7554/eLife.44532

4. Bauchle CJ, Rohli KE, Boyer CK, Pal V, Rocheleau JV, Liu S, Imai Y, Taylor EB, Stephens SB. 2021. Mitochondrial Efflux of Citrate and Isocitrate Is Fully Dispensable for Glucose-Stimulated Insulin Secretion and Pancreatic Islet β-Cell Function. Diabetes db210037. doi:10.2337/db21-0037

5. Bearrows SC, Bauchle CJ, Becker M, Haldeman JM, Swaminathan S, Stephens SB. 2019. Chromogranin B regulates early stage insulin granule trafficking from the Golgi in pancreatic islet beta-cells. J Cell Sci. doi:10.1242/jcs.231373

6. Birk J, Meyer M, Aller I, Hansen HG, Odermatt A, Dick TP, Meyer AJ, Appenzeller-Herzog C. 2013. Endoplasmic reticulum: reduced and oxidized glutathione revisited. J Cell Sci 126:1604–1617. doi:10.1242/jcs.117218

7. Boersema PJ, Raijmakers R, Lemeer S, Mohammed S, Heck AJ. 2009. Multiplex peptide stable isotope dimethyl labeling for quantitative proteomics. Nat Protoc 4:484–94. doi:10.1038/nprot.2009.21

8. Boland BB, Brown C, Alarcon C, Demozay D, Grimsby JS, Rhodes CJ. 2018. β-Cell Control of Insulin Production During Starvation-Refeeding in Male Rats. Endocrinology 159:895–906. doi:10.1210/en.2017-03120

9. Boland BB, Rhodes CJ, Grimsby JS. 2017. The dynamic plasticity of insulin production in β-cells. Mol Metab 6:958–973. doi:10.1016/j.molmet.2017.04.010

10. Cao M, Mao Z, Kam C, Xiao N, Cao X, Shen C, Cheng KKY, Xu A, Lee K-M, Jiang L, Xia J. 2013. PICK1 and ICA69 control insulin granule trafficking and their deficiencies lead to impaired glucose tolerance. PLoS Biol 11:e1001541. doi:10.1371/journal.pbio.1001541

11. Cao X, Lilla S, Cao Z, Pringle MA, Oka OBV, Robinson PJ, Szmaja T, van Lith M, Zanivan S, Bulleid NJ. 2020. The mammalian cytosolic thioredoxin reductase pathway acts via a membrane protein to reduce ER-localised proteins. J Cell Sci 133:jcs241976. doi:10.1242/jcs.241976

12. Delépine M, Nicolino M, Barrett T, Golamaully M, Lathrop GM, Julier C. 2000. EIF2AK3, encoding translation initiation factor 2-alpha kinase 3, is mutated in patients with Wolcott-Rallison syndrome. Nat Genet 25:406–409. doi:10.1038/78085

13. Ellgaard L, Sevier CS, Bulleid NJ. 2018. How Are Proteins Reduced in the Endoplasmic Reticulum? Trends Biochem Sci 43:32–43. doi:10.1016/j.tibs.2017.10.006

14. Engin F, Nguyen T, Yermalovich A, Hotamisligil GS. 2014. Aberrant islet unfolded protein response in type 2 diabetes. Sci Rep 4:4054. doi:10.1038/srep04054

15. Ferdaoussi M, Dai X, Jensen MV, Wang R, Peterson BS, Huang C, Ilkayeva O, Smith N, Miller N, Hajmrle C, Spigelman AF, Wright RC, Plummer G, Suzuki K, Mackay JP, van de Bunt M, Gloyn AL, Ryan TE, Norquay LD, Brosnan MJ, Trimmer JK, Rolph TP, Kibbey RG, Manning Fox JE, Colmers WF, Shirihai OS, Neufer PD, Yeh ETH, Newgard CB, MacDonald PE. 2015. Isocitrate-to-SENP1 signaling amplifies insulin secretion and rescues dysfunctional β cells. J Clin Invest 125:3847–3860. doi:10.1172/JCI82498

16. Gandasi NR, Yin P, Omar-Hmeadi M, Ottosson Laakso E, Vikman P, Barg S. 2018. Glucose-Dependent Granule Docking Limits Insulin Secretion and Is Decreased in Human Type 2 Diabetes. Cell Metab 27:470–478 e4. doi:10.1016/j.cmet.2017.12.017

17. Gansemer ER, McCommis KS, Martino M, King-McAlpin AQ, Potthoff MJ, Finck BN, Taylor EB, Rutkowski DT. 2020. NADPH and Glutathione Redox Link TCA Cycle Activity to Endoplasmic Reticulum Homeostasis. iScience 23:101116. doi:10.1016/j.isci.2020.101116

18. Gautier A, Juillerat A, Heinis C, Correa IR, Kindermann M, Beaufils F, Johnsson K. 2008. An engineered protein tag for multiprotein labeling in living cells. Chem Biol 15:128–36. doi:10.1016/j.chembiol.2008.01.007

19. Gembal M, Gilon P, Henquin JC. 1992. Evidence that glucose can control insulin release independently from its action on ATP-sensitive K+ channels in mouse B cells. J Clin Invest 89:1288–1295. doi:10.1172/JCI115714

20. Ghiasi SM, Dahlby T, Hede Andersen C, Haataja L, Petersen S, Omar-Hmeadi M, Yang M, Pihl C, Bresson SE, Khilji MS, Klindt K, Cheta O, Perone MJ, Tyrberg B, Prats C, Barg S, Tengholm A, Arvan P, Mandrup-Poulsen T, Marzec MT. 2019. Endoplasmic Reticulum Chaperone Glucose-Regulated Protein 94 Is Essential for Proinsulin Handling. Diabetes 68:747–760. doi:10.2337/db18-0671

21. Guest PC, Rhodes CJ, Hutton JC. 1989. Regulation of the biosynthesis of insulin-secretory-granule proteins. Co-ordinate translational control is exerted on some, but not all, granule matrix constituents. Biochem J 257:431–7.

22. Haataja L, Arunagiri A, Hassan A, Regan K, Tsai B, Dhayalan B, Weiss MA, Liu M, Arvan P. 2021. Distinct states of proinsulin misfolding in MIDY. Cell Mol Life Sci 78:6017–6031. doi:10.1007/s00018-021-03871-1

23. Halban PA, Polonsky KS, Bowden DW, Hawkins MA, Ling C, Mather KJ, Powers AC, Rhodes CJ, Sussel L, Weir GC. 2014. beta-cell failure in type 2 diabetes: postulated mechanisms and prospects for prevention and treatment. Diabetes Care 37:1751–8. doi:10.2337/dc14-0396

24. Han J, Song B, Kim J, Kodali VK, Pottekat A, Wang M, Hassler J, Wang S, Pennathur S, Back SH, Katze MG, Kaufman RJ. 2015. Antioxidants Complement the Requirement for Protein Chaperone Function to Maintain β-Cell Function and Glucose Homeostasis. Diabetes 64:2892–2904. doi:10.2337/db14-1357

25. Harding HP, Zeng H, Zhang Y, Jungries R, Chung P, Plesken H, Sabatini DD, Ron D. 2001. Diabetes mellitus and exocrine pancreatic dysfunction in perk-/- mice reveals a role for translational control in secretory cell survival. Mol Cell 7:1153–1163. doi:10.1016/s1097-2765(01)00264-7

26. Hassler JR, Scheuner DL, Wang S, Han J, Kodali VK, Li P, Nguyen J, George JS, Davis C, Wu SP, Bai Y, Sartor M, Cavalcoli J, Malhi H, Baudouin G, Zhang Y, Yates JR, Itkin-Ansari P, Volkmann N, Kaufman RJ. 2015. The IRE1α/XBP1s Pathway Is Essential for the Glucose Response and Protection of β Cells. PLoS Biol 13:e1002277. doi:10.1371/journal.pbio.1002277

27. Hayes HL, Peterson BS, Haldeman JM, Newgard CB, Hohmeier HE, Stephens SB. 2017. Delayed apoptosis allows islet beta-cells to implement an autophagic mechanism to promote cell survival. PLoS One 12:e0172567. doi:10.1371/journal.pone.0172567

28. Haythorne E, Rohm M, van de Bunt M, Brereton MF, Tarasov AI, Blacker TS, Sachse G, Silva Dos Santos M, Terron Exposito R, Davis S, Baba O, Fischer R, Duchen MR, Rorsman P, MacRae JI, Ashcroft FM. 2019. Diabetes causes marked inhibition of mitochondrial metabolism in pancreatic β-cells. Nat Commun 10:2474. doi:10.1038/s41467-019-10189-x

29. Henquin JC. 2009. Regulation of insulin secretion: a matter of phase control and amplitude modulation. Diabetologia 52:739–751. doi:10.1007/s00125-009-1314-y

30. Henquin JC, Ishiyama N, Nenquin M, Ravier MA, Jonas JC. 2002. Signals and pools underlying biphasic insulin secretion. Diabetes 51 **Suppl 1**:S60–7.

31. Hohmeier HE, Mulder H, Chen G, Henkel-Rieger R, Prentki M, Newgard CB. 2000. Isolation of INS-1-derived cell lines with robust ATP-sensitive K+ channel-dependent and - independent glucose-stimulated insulin secretion. Diabetes 49:424–30.

32. Hsu J-L, Huang S-Y, Chow N-H, Chen S-H. 2003. Stable-isotope dimethyl labeling for quantitative proteomics. Anal Chem 75:6843–6852. doi:10.1021/ac0348625

33. Hung V, Udeshi ND, Lam SS, Loh KH, Cox KJ, Pedram K, Carr SA, Ting AY. 2016. Spatially resolved proteomic mapping in living cells with the engineered peroxidase APEX2. Nat Protoc 11:456–75. doi:10.1038/nprot.2016.018

34. Ivarsson R, Quintens R, Dejonghe S, Tsukamoto K, in ’t Veld P, Renstrom E, Schuit FC. 2005. Redox control of exocytosis: regulatory role of NADPH, thioredoxin, and glutaredoxin. Diabetes 54:2132–42.

35. Jang I, Pottekat A, Poothong J, Yong J, Lagunas-Acosta J, Charbono A, Chen Z, Scheuner DL, Liu M, Itkin-Ansari P, Arvan P, Kaufman RJ. 2019. PDIA1/P4HB is required for efficient proinsulin maturation and ß cell health in response to diet induced obesity. Elife 8:e44528. doi:10.7554/eLife.44528

36. Joshi G, Chi Y, Huang Z, Wang Y. 2014. Aβ-induced Golgi fragmentation in Alzheimer’s disease enhances Aβ production. Proc Natl Acad Sci U S A 111:E1230–1239. doi:10.1073/pnas.1320192111

37. Kahn SE, Halban PA. 1997. Release of incompletely processed proinsulin is the cause of the disproportionate proinsulinemia of NIDDM. Diabetes 46:1725–32.

38. Kahn SE, Zraika S, Utzschneider KM, Hull RL. 2009. The beta cell lesion in type 2 diabetes: there has to be a primary functional abnormality. Diabetologia 52:1003–12. doi:10.1007/s00125-009-1321-z

39. Kang T, Boland BB, Alarcon C, Grimsby JS, Rhodes CJ, Larsen MR. 2019. Proteomic Analysis of Restored Insulin Production and Trafficking in Obese Diabetic Mouse Pancreatic Islets Following Euglycemia. J Proteome Res 18:3245–3258. doi:10.1021/acs.jproteome.9b00160

40. Kim D-S, Song L, Wang J, Wu H, Gu G, Sugi Y, Li Z, Wang H. 2018. GRP94 Is an Essential Regulator of Pancreatic β-Cell Development, Mass, and Function in Male Mice. Endocrinology 159:1062–1073. doi:10.1210/en.2017-00685

41. Lam SS, Martell JD, Kamer KJ, Deerinck TJ, Ellisman MH, Mootha VK, Ting AY. 2015. Directed evolution of APEX2 for electron microscopy and proximity labeling. Nat Methods 12:51– 4. doi:10.1038/nmeth.3179

42. Like AA, Chick WL. 1970. Studies in the diabetic mutant mouse. II. Electron microscopy of pancreatic islets. Diabetologia 6:216–242. doi:10.1007/BF01212232

43. Lipson KL, Fonseca SG, Ishigaki S, Nguyen LX, Foss E, Bortell R, Rossini AA, Urano F. 2006. Regulation of insulin biosynthesis in pancreatic beta cells by an endoplasmic reticulum-resident protein kinase IRE1. Cell Metab 4:245–254. doi:10.1016/j.cmet.2006.07.007

44. Liu M, Hodish I, Rhodes CJ, Arvan P. 2007. Proinsulin maturation, misfolding, and proteotoxicity. Proc Natl Acad Sci U S A 104:15841–6. doi:10.1073/pnas.0702697104

45. Malhotra JD, Miao H, Zhang K, Wolfson A, Pennathur S, Pipe SW, Kaufman RJ. 2008. Antioxidants reduce endoplasmic reticulum stress and improve protein secretion. Proc Natl Acad Sci U S A 105:18525–18530. doi:10.1073/pnas.0809677105

46. Marchetti P, Bugliani M, Lupi R, Marselli L, Masini M, Boggi U, Filipponi F, Weir GC, Eizirik DL, Cnop M. 2007. The endoplasmic reticulum in pancreatic beta cells of type 2 diabetes patients. Diabetologia 50:2486–2494. doi:10.1007/s00125-007-0816-8

47. Masini M, Marselli L, Bugliani M, Martino L, Masiello P, Marchetti P, De Tata V. 2012. Ultrastructural morphometric analysis of insulin secretory granules in human type 2 diabetes. Acta Diabetol 49 **Suppl 1**:S247–252. doi:10.1007/s00592-012-0446-6

48. Muller A, Neukam M, Ivanova A, Sonmez A, Munster C, Kretschmar S, Kalaidzidis Y, Kurth T, Verbavatz JM, Solimena M. 2017. A Global Approach for Quantitative Super Resolution and Electron Microscopy on Cryo and Epoxy Sections Using Self-labeling Protein Tags. Sci Rep 7:23. doi:10.1038/s41598-017-00033-x

49. Ozcan U, Yilmaz E, Ozcan L, Furuhashi M, Vaillancourt E, Smith RO, Görgün CZ, Hotamisligil GS. 2006. Chemical chaperones reduce ER stress and restore glucose homeostasis in a mouse model of type 2 diabetes. Science 313:1137–1140. doi:10.1126/science.1128294

50. Pasquier A, Vivot K, Erbs E, Spiegelhalter C, Zhang Z, Aubert V, Liu Z, Senkara M, Maillard E, Pinget M, Kerr-Conte J, Pattou F, Marciniak G, Ganzhorn A, Ronchi P, Schieber NL, Schwab Y, Saftig P, Goginashvili A, Ricci R. 2019. Lysosomal degradation of newly formed insulin granules contributes to β cell failure in diabetes. Nat Commun 10:3312. doi:10.1038/s41467-019-11170-4

51. Perez-Riverol Y, Csordas A, Bai J, Bernal-Llinares M, Hewapathirana S, Kundu DJ, Inuganti A, Griss J, Mayer G, Eisenacher M, Pérez E, Uszkoreit J, Pfeuffer J, Sachsenberg T, Yilmaz S, Tiwary S, Cox J, Audain E, Walzer M, Jarnuczak AF, Ternent T, Brazma A, Vizcaíno JA. 2019. The PRIDE database and related tools and resources in 2019: improving support for quantification data. Nucleic Acids Res 47:D442–D450. doi:10.1093/nar/gky1106

52. Reinbothe TM, Ivarsson R, Li D-Q, Niazi O, Jing X, Zhang E, Stenson L, Bryborn U, Renström E. 2009. Glutaredoxin-1 Mediates NADPH-Dependent Stimulation of Calcium-Dependent Insulin Secretion. Molecular Endocrinology 23:893–900. doi:10.1210/me.2008-0306

53. Reissaus CA, Piñeros AR, Twigg AN, Orr KS, Conteh AM, Martinez MM, Kamocka MM, Day RN, Tersey SA, Mirmira RG, Dunn KW, Linnemann AK. 2019. A Versatile, Portable Intravital Microscopy Platform for Studying Beta-cell Biology In Vivo. Sci Rep 9:8449. doi:10.1038/s41598-019-44777-0

54. Rhodes CJ, Alarcon C. 1994. What beta-cell defect could lead to hyperproinsulinemia in NIDDM? Some clues from recent advances made in understanding the proinsulin-processing mechanism. Diabetes 43:511–7.

55. Ronnebaum SM, Ilkayeva O, Burgess SC, Joseph JW, Lu D, Stevens RD, Becker TC, Sherry AD, Newgard CB, Jensen MV. 2006. A pyruvate cycling pathway involving cytosolic NADP-dependent isocitrate dehydrogenase regulates glucose-stimulated insulin secretion. J Biol Chem 281:30593–602. doi:M511908200 [pii] 10.1074/jbc.M511908200

56. Sharma RB, O’Donnell AC, Stamateris RE, Ha B, McCloskey KM, Reynolds PR, Arvan P, Alonso LC. 2015. Insulin demand regulates β cell number via the unfolded protein response. J Clin Invest 125:3831–3846. doi:10.1172/JCI79264

57. Shrestha N, Liu T, Ji Y, Reinert RB, Torres M, Li X, Zhang M, Tang C-HA, Hu C-CA, Liu C, Naji A, Liu M, Lin JD, Kersten S, Arvan P, Qi L. 2020. Sel1L-Hrd1 ER-associated degradation maintains β cell identity via TGF-β signaling. J Clin Invest 130:3499–3510. doi:10.1172/JCI134874

58. Stephens SB, Edwards RJ, Sadahiro M, Lin WJ, Jiang C, Salton SR, Newgard CB. 2017. The Prohormone VGF Regulates beta Cell Function via Insulin Secretory Granule Biogenesis. Cell Rep 20:2480–2489. doi:10.1016/j.celrep.2017.08.050

59. Støy J, Steiner DF, Park S-Y, Ye H, Philipson LH, Bell GI. 2010. Clinical and molecular genetics of neonatal diabetes due to mutations in the insulin gene. Rev Endocr Metab Disord 11:205–215. doi:10.1007/s11154-010-9151-3

60. Tran DT, Pottekat A, Mir SA, Loguercio S, Jang I, Campos AR, Scully KM, Lahmy R, Liu M, Arvan P, Balch WE, Kaufman RJ, Itkin-Ansari P. 2020. Unbiased Profiling of the Human Proinsulin Biosynthetic Interaction Network Reveals a Role for Peroxiredoxin 4 in Proinsulin Folding. Diabetes 69:1723–1734. doi:10.2337/db20-0245

61. Uchizono Y, Alarcón C, Wicksteed BL, Marsh BJ, Rhodes CJ. 2007. The balance between proinsulin biosynthesis and insulin secretion: where can imbalance lead? Diabetes Obes Metab 9 **Suppl 2**:56–66. doi:10.1111/j.1463-1326.2007.00774.x

62. Ward WK, Bolgiano DC, McKnight B, Halter JB, Porte D. 1984. Diminished B cell secretory capacity in patients with noninsulin-dependent diabetes mellitus. J Clin Invest 74:1318– 28. doi:10.1172/JCI111542

63. Wicksteed B, Alarcon C, Briaud I, Lingohr MK, Rhodes CJ. 2003. Glucose-induced translational control of proinsulin biosynthesis is proportional to preproinsulin mRNA levels in islet beta-cells but not regulated via a positive feedback of secreted insulin. J Biol Chem 278:42080–42090. doi:10.1074/jbc.M303509200

64. Winter J, Klappa P, Freedman RB, Lilie H, Rudolph R. 2002. Catalytic activity and chaperone function of human protein-disulfide isomerase are required for the efficient refolding of proinsulin. J Biol Chem 277:310–317. doi:10.1074/jbc.M107832200

65. Xin Y, Dominguez Gutierrez G, Okamoto H, Kim J, Lee A-H, Adler C, Ni M, Yancopoulos GD, Murphy AJ, Gromada J. 2018. Pseudotime Ordering of Single Human β-Cells Reveals States of Insulin Production and Unfolded Protein Response. Diabetes 67:1783–1794. doi:10.2337/db18-0365

66. Xu T, Yang L, Yan C, Wang Xiaoxia, Huang P, Zhao F, Zhao L, Zhang M, Jia W, Wang Xiangdong, Liu Y. 2014. The IRE1α-XBP1 pathway regulates metabolic stress-induced compensatory proliferation of pancreatic β-cells. Cell Res 24:1137–1140. doi:10.1038/cr.2014.55

67. Yong J, Johnson JD, Arvan P, Han J, Kaufman RJ. 2021. Therapeutic opportunities for pancreatic β-cell ER stress in diabetes mellitus. Nat Rev Endocrinol 17:455–467. doi:10.1038/s41574-021-00510-4

68. Yu CL, Brooks S, Li Y, Subramanian M, Summers R, Pope M. 2017. Rapid Proteomics to Prospect and Validate Novel Bacterial Metabolism Induced by Environmental Burden. Methods Enzymol 586:379–411. doi:10.1016/bs.mie.2016.11.003

69. Zougman A, Selby PJ, Banks RE. 2014. Suspension trapping (STrap) sample preparation method for bottom-up proteomics analysis. Proteomics 14:1006–1000. doi:10.1002/pmic.201300553

